# spCorr: flexible and scalable inference of spatially varying correlation in spatial transcriptomics

**DOI:** 10.1101/2025.09.30.679684

**Authors:** Chenxin Flora Jiang, Yuxin Yin, Paul Robson, Yuanhao James Li, Jingyi Jessica Li, Dongyuan Song

## Abstract

Spatial transcriptomics has transformed our ability to explore gene expression within its tissue context, enabling us to dissect subtle yet biologically significant variations *in situ*. While numerous computational methods have been proposed for detecting Spatially Varying Genes (SVGs) expression by modeling each individual gene separately, much less effort has been devoted to understanding how correlations between genes change across space. Such Spatially Varying Correlations (SVCs) are critical for understanding biological processes such as gene regulatory mechanisms shaped by local tissue environments, yet existing tools remain limited for this task. To address this gap, we present spCorr, a flexible and scalable regression framework for studying SVCs. spCorr provides interpretable, spot-level estimates of gene correlation and detects gene pairs whose correlations vary across locations or between tissue domains. Through extensive simulations and real-data analyses, we show that spCorr achieves high detection power, reliably controls the False Discovery Rate (FDR), and is computationally efficient. Importantly, spCorr reveals biologically meaningful correlation patterns that highlight fine-scale tissue structures, gene module functions, and region-specific interactions, offering new opportunities to study coordinated gene regulation in spatial transcriptomics.

## 1 Introduction

Understanding how genes coordinate their activity across biological contexts has led to rich insights into biological processes and disease mechanisms. Genes do not function in isolation; instead, genes operate within coordinated networks that dynamically adapt across tissues, spatial locations, developmental stages, environmental conditions, and disease states [1–4]. Over the past two decades, gene correlation studies have become increasingly crucial for exploring the system-level functionality of genes, supported by technologies such as microarrays and RNA sequencing [5–7]. While in literature, the terms “gene correlation” and “gene co-expression” are often used interchangeably, we use “gene correlation” in this manuscript to represent continuous variation rather than the binary “expressed or not” case. Most existing computational methods for analyzing gene correlation are developed for bulk RNA sequencing or single-cell RNA sequencing (scRNA-seq). Bulk approaches capture gene correlations across entire tissues [8], while scRNA-seq enables the estimation of correlations within specific cell types [9, 10]. Although these approaches provide valuable insight into gene regulatory mechanisms at the population level, they are limited to aggregated views based on grouped cells or cell subpopulations and cannot reflect spatially localized variation in gene correlation within fine-scale tissue structures.

Recent advances in spatial transcriptomics (ST) technologies offer an opportunity to study gene correlation in a spatially resolved context. By retaining spatial information alongside gene expression, both sequencing-based [11–13] and imaging-based [14–16] ST platforms are enabling new insights into how the local tissue environment shapes gene activity [17]. Different ST technologies may have different resolutions, from multi-cell to single-cell. In this paper, we refer to the smallest unit, either cell or captured binned square, as “spot”. To date, a primary focus of ST data analysis has been the identification of spatially variable genes (SVGs), that is, genes that exhibit significant expression variation across spatial locations. More than 30 computational methods have been developed for this task [18]. However, these methods focus exclusively on changes in the marginal expression of individual genes and do not consider how gene-gene relationships may vary across space. This limitation is not merely computational; it risks overlooking key biological phenomena. Spatially localized co-regulation, driven by cell-cell interactions, tissue architecture, and the microenvironment, is fundamental to tissue function. Indeed, a growing body of evidence shows that spatial location strongly influences coordinated gene expression and that these coordinated changes are implicated in development, disease, and therapeutic response [19–21]. These observations highlight the need for computational methods capable of detecting spatial variation in gene correlation from ST data. By analogy to SVG identification, we term this complementary and critical task the identification of Spatially Varying Correlation (**SVC**).

To our knowledge, only a few methods are specifically designed for identifying SVCs in ST data. These methods generally follow a two-step, non-parametric approach: (i) local correlation estimation and (ii) hypothesis testing for SVC. For the first step, methods estimate the correlation between gene pairs at each spot. For example, SpatialDM [22] uses a bivariate Moran’s statistic with predefined spatial weights, while methods like scHOT [23] and SpatialCorr [24] use kernel-based weighting to estimate local Spearman or Pearson correlations. For the second step, these methods identify SVC by evaluating whether the estimated local correlations vary significantly across space. This is typically achieved through permutation-based testing or, in the case of SpatialDM, by approximating the null distribution with an analytic derivation. While these methods established the feasibility of SVC analysis, they have several key limitations. First, their correlation estimates are difficult to interpret, as they cannot be expressed as explicit functions of spatial coordinates. Second, they lack a framework for adjusting for extra spot-level covariates associated with gene expression, such as library size or domain differences, which may confound the estimation of the interested correlation. Third, they are not designed to directly model count data, often requiring transformation procedures that can distort correlation structures. Finally, their reliance on permutation tests is computationally expensive, while their analytical assumptions can be restrictive, potentially leading to inaccurate *p*-values.

To address these challenges, we introduce spCorr, the first interpretable, regression-based method for identifying spatially varying correlation and inferring local gene correlation in ST data. By building a novel regression model based on the product of two genes’ transformed expression values, spCorr offers several key advantages: (1) it estimates local correlations as an explicit, interpretable function of spatial locations; (2) it allows for flexible adjustment of spot-level covariates to prevent confounding; (3) it directly models count data, avoiding potentially distortion from transformations; and (4) it provides computationally efficient and statistically rigorous hypothesis testing without relying on permutations. A summary comparison of spCorr with existing methods is provided in Table S1.

The spCorr method shows strong performance in both simulation studies and real data applications. In simulations using synthetic and semi-synthetic data with known ground truth, spCorr was benchmarked against SpatialDM, scHOT, and SpatialCorr [22–24]. It is the only method that effectively controls the false discovery rate (FDR) while also achieving the highest power in SVC identification. In addition, spCorr exhibits superior computational efficiency compared to other methods. To showcase its practical utility, we applied spCorr to three ST datasets generated by three mainstream ST technologies, which cover both sequencing-based and imaging-based platforms, including the human oral squamous cell carcinoma dataset from the 10x Visium platform, the mouse cortex dataset from the 10x Xenium platform, and the mouse hippocampus dataset from the 10x Visium HD platform. In each case, spCorr identifies SVCs associated with biologically meaningful functions, either across a continuous spatial space or between discrete spatial domains. Furthermore, clustering based on spCorr’s local correlation estimates reveals subtle spatial structures, such as intratumoral architecture and brain cortical areas, which are invisible in clustering based on gene expression. Finally, applying spCorr to differential correlation network analysis uncovers gene-module-level discrepancies between the two brain regions.

## 2 Results

### 2.1 Overview of spCorr method

The spCorr offers a flexible regression framework to analyze how gene correlations vary across space, while adjusting for extra covariates that may confound correlation estimates. The required input of spCorr is shown in Fig. 1, left. Specifically, we denote the spot-by-gene count matrix as

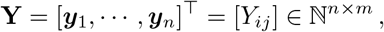

where *n* is the number of spots and *m* is the number of genes. Here we refer to capture locations, regardless of the single cell resolution or not, as “spots” for brevity, thus not restricting spCorr to experimental platforms or resolutions. Each spot has a spatial coordinate *s*_*i*_ = (*s*_*i*1_, *s*_*i*2_)^⊤^, collected in the matrix

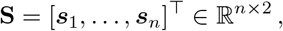

and may also have spot-level covariates (e.g., library size, spot annotation) stored in

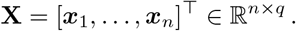

**Fig. 1:**
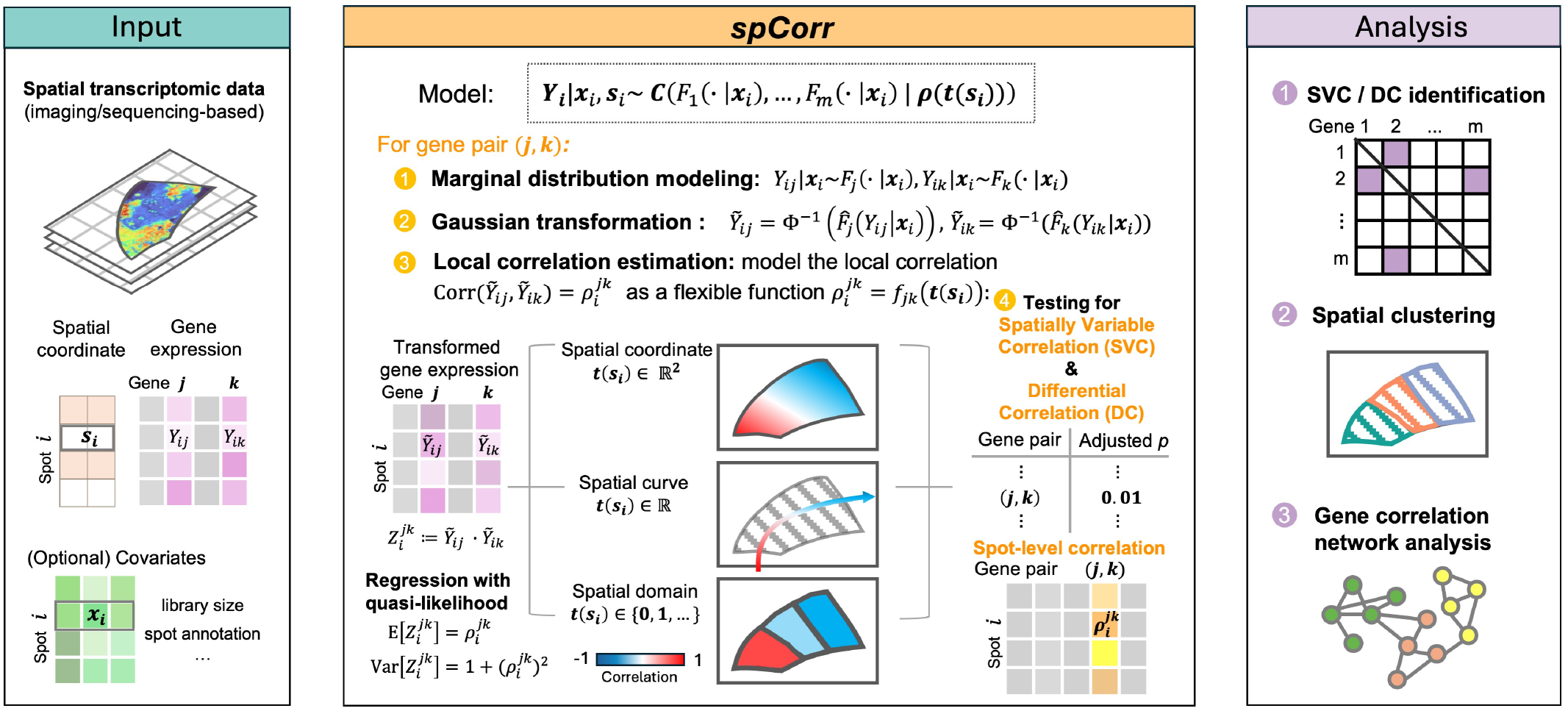
Overview of the spCorr framework for modeling spatially varying gene correlation in spatial transcriptomics (ST) data. spCorr is a statistical method that models how gene correlation varies across spatial locations in ST data. Input: The method takes as input spatial transcriptomics data, including spatial coordinates, gene expression levels, and optional spot-level covariates (e.g., library size and spot annotations) that may confound correlation estimates. Modeling: For each gene pair, spCorr performs four main steps: (1) marginal distribution modeling using generalized linear models (GLMs) to account for the confounding effects from spot-level covariates; (2) Gaussian transformation of gene expression values via probability integral transform and inverse normal transformation to standard Gaussian variables; (3) local correlation estimation by modeling the cross-product of transformed expressions of gene pairs using a quasi-likelihood generalized additive model (Quasi-GAM), using spatial coordinates, curves, or domain label as spatial predictors; and (4) hypothesis testing to identify (i) spatially varying correlations (SVCs) across continuous space and (ii) differential correlations (DCs) across predefined spatial domains. Output and analysis: spCorr yields interpretable spot-level correlation estimates and supports downstream tasks such as SVC/DC identification, spatial clustering based on correlation structure, and gene correlation network analysis.

The goal is to estimate spot-level, spatially-aware gene correlations. Since the dependency structure of count distributions (e.g., multivariate negative binomial) may not be well-defined, spCorr employs a Gaussian-copula model to estimate gene-gene dependence without relying on multivariate count distributions (Fig. 1, middle). Specifically, the expression distribution of each gene *j* is modeled conditionally on covariates ***x***_*i*_, written as *F*_*j*_(· | ***x***_*i*_). These marginal models capture gene-specific behaviors, while the correlation structure between genes is linked by a Gaussian copula. The correlation structure is parameterized by a spatially varying correlation matrix ***ρ***(***t***(***s***_*i*_)), whose entries represent the local correlation between gene pairs at spot *i*, where ***t***(***s***_*i*_) are the spatial predictors encoding different types of spatial information, such as two-dimensional spatial coordinates, a one-dimensional spatial curve, or discrete spatial domain labels associated with spot *i*. Although our previous work scDesign3 [25] has employed Gaussian copulas to model high-dimensional spatial transcriptomics data, it assumed the gene-gene correlation matrix was constant in space. spCorr overcomes this limitation by allowing the correlation structure to vary flexibly with ***t***(***s***_*i*_), thus adapting to diverse spatial contexts.

Estimating the high-dimensional correlation matrix is computationally challenging and often unnecessary, since researchers are typically interested in specific gene pairs (e.g., transcription factor - target, ligand - receptor, etc.). Therefore, spCorr adopts a pairwise strategy instead of estimating the entire high-dimensional correlation matrix. For a gene pair (*j, k*), the local correlation at spot *i* is assumed to be

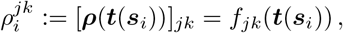

where *f*_*jk*_ describes how the correlation between genes *j* and *k* changes with spatial context.

To accurately estimate the local correlation 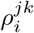, spCorr consists of four major steps: marginal distribution modeling, Gaussian transformation, local correlation estimation, and hypothesis testing (Fig. 1, middle). Given a target gene pair (*j, k*), spCorr performs the following four steps sequentially:

1. **Marginal distribution modeling:** For each gene pair, the expression values *Y*_*ij*_ and *Y*_*ik*_ across spots are modeled conditional on spot-level covariates ***x***_*i*_ using a generalized linear model (GLM). This step accounts for confounding variation (e.g., library size, spot annotation) that may affect the marginal distribution prior to correlation analysis. It is conceptually similar to the single-cell normalization method sctransform [26], but allows for more flexible specification.
2. **Gaussian transformation:** The fitted marginal models *Y*_*ij*_ | ***x***_*i*_ and *Y*_*ik*_ | ***x***_*i*_ are then used to transform the observed expression values into standard Gaussian variables 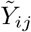 and 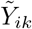. By mapping any gene’s expression quantiles to the standard Gaussian scale, this step places all genes on a unified scale suitable for correlation modeling.
3. **Local correlation estimation:** For each spot *i*, spCorr defines the cross-product between genes *j* and *k* as

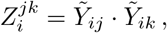

which has moment properties (its expectation and variance) directly related to local correlation *ρ*^*jk*^. By modeling *Z*_*i*_ as a function of the spatial predictor ***t***(***s***_*i*_) using a quasi-likelihood generalized additive model, spCorr obtains estimates of the spot-level correlation as an interpretable function of spatial locations

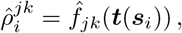

where *f*_*jk*_ can be chosen flexibly based on biological contexts.
4. **Hypothesis testing:** Finally, spCorr tests if the estimated local correlation varies significantly in space or not. spCorr supports two complementary types of test: (i) testing *spatially varying correlation (SVC)*, which evaluates whether correlations change continuously across spatial coordinates or spatial curves; and (ii) testing *differential correlation (DC)*, which compares correlations between discrete spatial domains.

Downstream analyses supported by spCorr include identifying gene pairs with SVC/DC, spatial clustering based on the spot-level correlation matrix, and gene network analysis (Fig. 1, right). Further details of spCorr are provided in Section 4.

### 2.2 Simulations verify spCorr’s reliable FDR control, high power, and computational efficiency

We conducted extensive simulation studies to validate spCorr as an effective method for identifying SVC in ST data, demonstrating reliable FDR control, high statistical power, and computational efficiency. In simulation setting 1, to ensure biological realism, we adopted a semi-synthetic simulation framework based on human dorsolateral prefrontal cortex (DLPFC) 10x Genomics Visium data [27]. From the top 400 spatially variable genes, we selected 200 gene pairs and generated synthetic gene expression using a bivariate Poisson–lognormal model, explicitly dividing them into SVC (spatially varying correlation) and non-SVC (constant correlation) groups. The simulation workflow (Fig. S1) combined spot-level library sizes, region-specific gene means, and Poisson sampling to mimic real ST data, with each setting replicated 30 times for robust evaluation. Full details of the simulation design in simulation setting 1 are provided in Supplementary Method S1.1.

We applied spCorr and three other methods developed for SVC identification and gene local correlation estimation in ST—scHOT, SpatialDM, and SpatialCorr [22–24]—to the simulated data. Our first evaluation assessed the validity of these methods’ *p*-values, which, under the null hypothesis (i.e., this gene pair is not SVC), are expected to follow a uniform distribution between 0 and 1. Our results indicate that, among the four methods, spCorr produces the best-calibrated *p*-values, closely matching the expected uniform distribution (Fig. 2a). Among the remaining methods, scHOT and SpatialCorr exhibit substantial deviation from the expected uniform distribution, while SpatialDM is completely off with an incorrect concentration of small *p*-values.

**Fig. 2:**
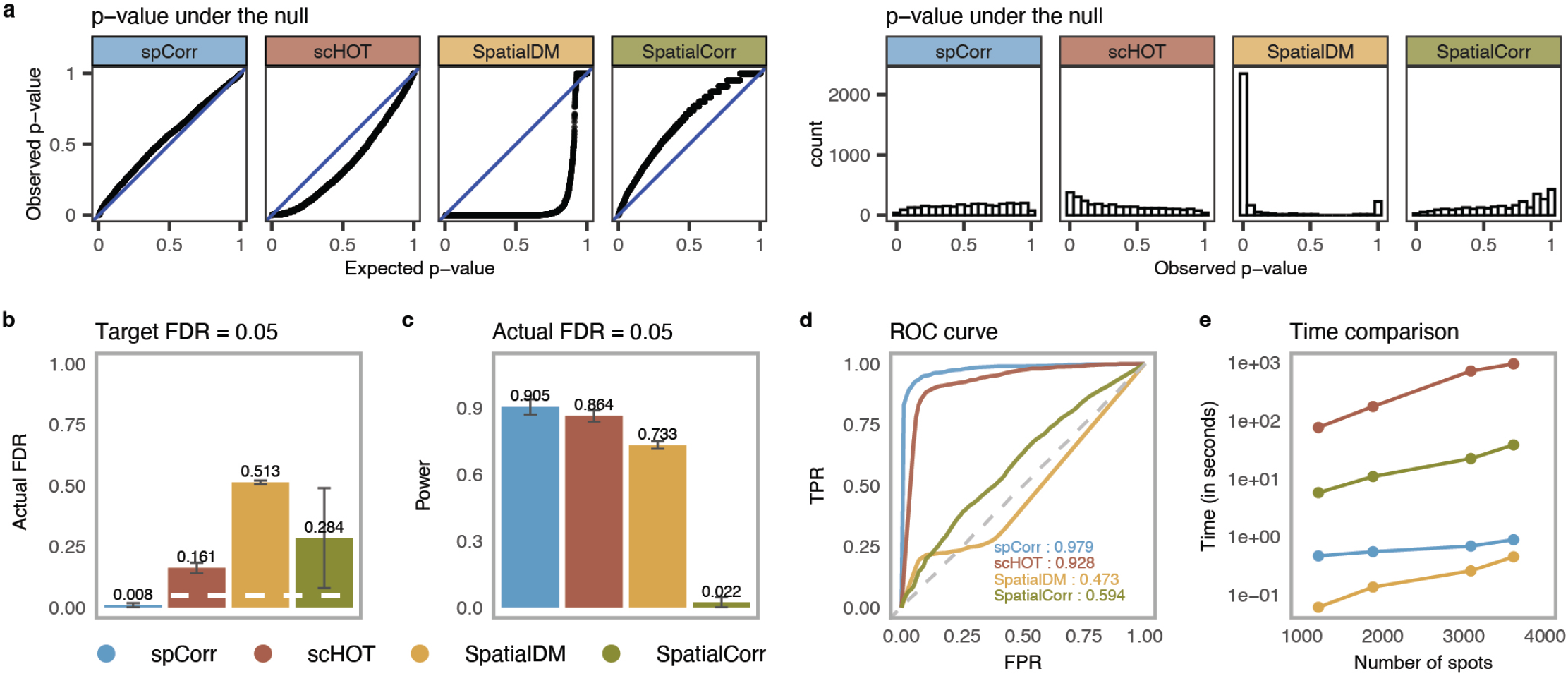
spCorr achieves reliable FDR control, high statistical power, and computational efficiency in simulation studies. **a**. Distributions of observed *p*-values for non-SVC gene pairs across four methods: spCorr, scHOT, SpatialDM, and SpatialCorr. Left: quantile-quantile plots comparing empirical *p*-value quantiles to the expected Uniform[0,1] distribution. Right: histograms of observed *p*-values. spCorr exhibits the closest adherence to the expected Uniform[0,1] distribution, indicating well-calibrated *p*-values. **b**. Actual FDRs achieved by each method under a target FDR of 0.05 (Benjamini-Hochberg adjusted *p* ≤ 0.05). spCorr shows the most accurate FDR control, while all other methods exceed the target threshold. **c**. Statistical power of the four methods under the actual FDR = 0.05 cutoff. spCorr achieves the highest power. **d**. ROC curves and corresponding AUROC values for the four methods. spCorr attains the highest AUROC among all methods. **e**. Runtime (in seconds per gene pair) versus the number of spots. spCorr demonstrates substantially better computational scalability than scHOT and SpatialCorr.

Our second evaluation focused on FDR control and the power of spCorr compared to the other methods. At a target 5% FDR threshold, spCorr is the only method that effectively controls the false discovery rate, whereas all three alternative methods fail to maintain FDR within the target threshold (Fig. 2b). When comparing statistical power under the 5% FDR threshold, spCorr achieves the highest power (Fig. 2c). We further assessed each method’s ability to distinguish SVCs from non-SVCs using the area under the receiver operating characteristic curve (AUROC) as the evaluation metric (Fig. 2d). Our results show that spCorr achieves the highest AUROC values, followed closely by scHOT, while SpatialDM and SpatialCorr show substantially lower performance. These findings highlight spCorr’s superior power and its ability to effectively control the FDR in identifying SVCs. To assess the impact of covariate adjustment in marginal distribution modeling (step 1), we compared the default version of spCorr (with covariate adjustment) against spCorr (without covariate adjustment). The result show that including covariates led to improved FDR control, higher power, and higher AUROC (Fig. S3), underscoring the importance of covariate adjustment in step 1 of the spCorr framework.

Another key advantage of spCorr is its computational efficiency. In our third evaluation, we examined the running time of each method across varying numbers of spatial spots. For each setting, we recorded the average running time per gene pair (Fig. 2e). Among the methods compared, SpatialDM exhibits the fastest runtime, followed by spCorr, which shows a substantial speed advantage over both SpatialCorr and scHOT. As a supplement to the semi-synthetic simulation, we also conducted a fully synthetic simulation. In simulation setting 2, we simulated data based on a bivariate negative binomial model with spatially varying copula structure. Again, spCorr maintained accurate FDR control and achieved the highest power and AUROC (Fig. S5). In summary, across diverse simulation settings, spCorr outperformed existing methods in SVC detection accuracy while maintaining strong computational efficiency.

### 2.3 spCorr deciphers heterogeneous regulatory mechanisms within oral squamous cell carcinoma

In our first real-data application, we applied spCorr to ST data of HPV-negative oral squamous cell carcinoma (OSCC) from the 10x Genomics Visium platform [21]. We used slice 2 as the example, which contains expression measurements for 15,624 genes across 1,749 spots. Pathologists divided the Hematoxylin and Eosin (H&E)-stained histological image into six tissue categories, including one tumor region (Tumor) and four non-tumor regions(Fig. 3a). To adjust for potential expression differences between tumor and non-tumor tissues, we included a binary covariate indicating tumor presence in step 1 of spCorr. This helped deconfound correlation estimates and focus the analysis on intratumoral variation. To reduce the search space and focus on biologically relevant interactions, we selected transcription factor (TF)-target gene pairs from the TRRUST v2 database [28]. We further refined this set by removing those regulatory interactions classified as “Repression” and filtered out genes with expression in fewer than 10% of spots.

**Fig. 3:**
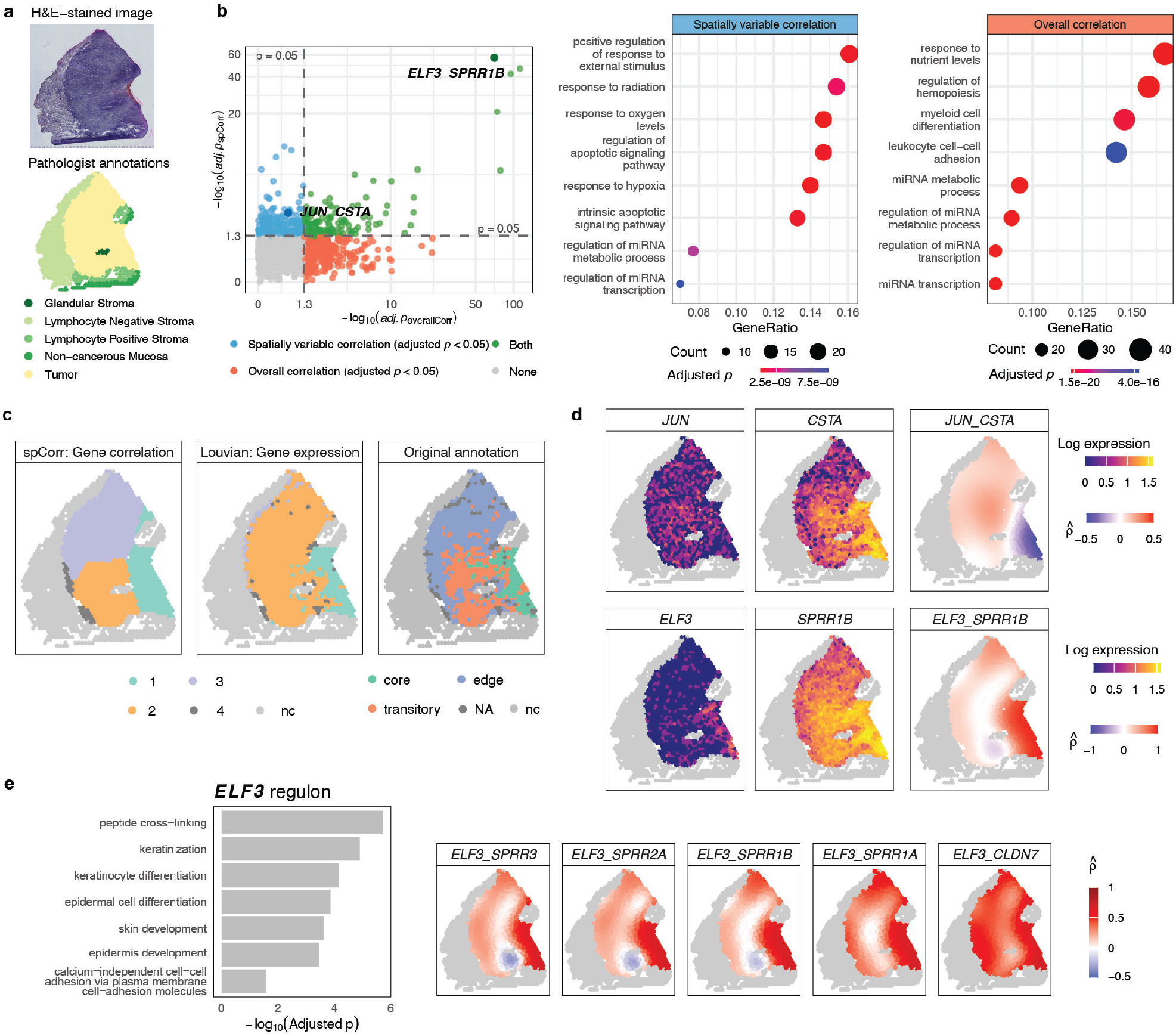
spCorr elucidates intratumoral regulatory heterogeneity. **a**. H&E-stained image and pathologist-defined spatial annotations for sample 2 of the HPV-negative OSCC dataset. The tumor region is distinguished from the surrounding stromal and mucosal tissue. **b**. Comparison of spatially varying correlation (SVC) versus non-spatial overall correlation across all candidate TF-target gene pairs. Left: overlap of gene pairs identified as significant (Benjamini-Hochberg adjusted *p <* 0.05) by either or both approaches. Right: GO enrichment analysis shows that genes from SVC pairs are enriched in terms related to tumor progression. **c**. Left: Spatial clustering based on spCorr-inferred local correlation estimates reveals tumor subdomains that aligned with original annotations (right) while offering smoother and more clearly defined boundaries. Middle: Louvain clustering based on expression of the same TF-target gene pairs fails to produce comparably distinct boundaries. **d**. Spot-level correlation estimates for two representative TF-target pairs. Top: the *JUN* and *CSTA* pair exhibits a negative correlation in the tumor core and a positive correlation in the leading edge and transitory region, suggesting localized inflammatory activity in tumor peripheral areas. Bottom: the *ELF3* and *SPRR1B* pair displays a strong positive correlation in the tumor core, consistent with *ELF3* ‘s known role in epithelial differentiation. **e**. Functional analysis of the *ELF3* regulon. Left: GO enrichment analysis of *ELF3* regulon reveals biological processes including keratinization and epidermal development. Right: local correlation estimates for *ELF3* and its targets are aligned with the boundary of the tumor core, indicating their domain-specific transcriptional activities.

To assess the utility of spCorr in detecting spatially varying transcriptional regulation, we first compared its output to a standard, non-spatial correlation analysis. When applied to the 1,316 candidate TF-target gene pairs, spCorr identified 334 TF-target pairs exhibiting significant SVC at the 5% FDR threshold. For comparison, we also computed the overall correlation (non-spatial, naïve Pearson correlation) and corresponding *p*-value for each gene pair, identifying 417 significantly correlated pairs at the same FDR threshold (Fig. 3b, left). These two types of correlation show a clear discrepancy in their significance pairs (Fig. 3b). To further evaluate the biological relevance of the identified pairs, we performed Gene Ontology (GO) functional enrichment analysis for genes identified through spatially varying correlations and those identified through overall correlations, respectively. Genes involved in spatially varying correlations were significantly enriched for biological processes commonly associated with solid tumor progression, including the positive regulation of responses to external stimuli, oxygen levels, and hypoxia-hallmark features [29]. In contrast, genes identified solely based on overall correlation were enriched in functions barely related to tumor biology, highlighting the added biological insight gained through spatially resolved correlation analysis (Fig. 3b, right).

Next, we explored whether local correlation patterns could reveal spatial heterogeneity within the tumor region. We clustered the spots based on their spCorr-inferred local correlation estimates from the significant SVC pairs. This correlation-based clustering revealed four distinct spatial domains (Fig. 3c, left) that closely aligned with the annotations reported in the original paper based on multi-sample integration (Fig. 3c, right). Specifically, clusters 1, 2, and 3 derived from spCorr strongly corresponded to the original annotations of the tumor core, transitory region, and leading edge, respectively, while offering smoother patterns and more clearly delineated spatial boundaries. Remarkably, spCorr achieved this refined clustering using only a small subset of genes from significant TF-target pairs based on one sample (sample 2), whereas the original annotations relied on whole transcriptomics across twelve samples. In contrast, Louvain clustering based solely on gene expression profiles of the same TF-target pairs resulted in a less distinct spatial pattern (Fig. 3c, middle).

To gain mechanistic insights into local regulatory programs, we investigated local correlation patterns for specific TF-target pairs from spCorr. The pair *JUN* (c-Jun) and *CSTA* (Cystatin A), for example, exhibited a clear spatial shift in correlation pattern, with a negative correlation in the tumor core that became positive in the leading edge (Fig. 3d, top). This pattern suggests localized activation of inflammatory programs in the peripheral areas of the tumor, as *JUN*, a key member of the AP-1 complex, has been shown to drive inflammation through increased pro-inflammatory signaling [30]. Another pair, *ELF3* (E74-like factor 3) and *SPRR1B* (Small proline-rich protein 1B), showed a strong positive correlation restricted to the tumor core (Fig. 3d, bottom). This may reflect active epithelial differentiation programs, as *ELF3* is known to regulate *SPRR1B* expression in keratinocytes and plays a key role in cornification and epithelial barrier formation [31]. To further characterize the regulatory role of *ELF3*, we extended our analysis from individual gene pairs to its broader set of target genes. Using the spot-level correlation estimates for each TF-target pair, we identified the potential downstream target genes of *ELF3*, thereby constructing its regulon. GO analysis of the *ELF3* regulon showed significant enrichment for biological processes such as keratinization, keratinocyte differentiation, and epidermal cell differentiation (Fig. 3e, left), consistent with previously reported functions of *ELF3* [32]. Notably, the spatial pattern of correlation between *ELF3* and its targets closely aligned with the tumor core boundary (Fig. 3e, right), suggesting domain-specific activation of *ELF3* -regulated transcriptional programs.

### 2.4 spCorr uncovers mouse brain cortex functional regions with a limited number of genes

In the second real data application, we applied spCorr to ST data of a coronal section of the mouse brain generated by the 10x Genomics Xenium In Situ technology. Unlike sequencing-based technologies, which measure the whole transcriptome (e.g., 10x Visium in the first real data application), Xenium technology only measures a subset of genes from the pre-defined panel, which is 248 genes in this dataset. We focused on studying the cortex structure, thus subsetting a cortex region with 7, 477 cells. We used two supervised cell-type annotation algorithms to obtain faithful cell-type labels (Fig. 4a, left). The cell type labels, particularly those corresponding to cortical neurons, clearly align with the six layers of the cerebral cortex (Fig. 4a, right; Allen reference as the ground truth). However, it is easy to notice that the functional region segmentations that are horizontal to the cerebral cortex, e.g., Retrosplenial Cortex (RSP), Visual Cortex (VIS), and Primary Somatosensory Cortex (SSp), are not reflected by cell type layers.

**Fig. 4:**
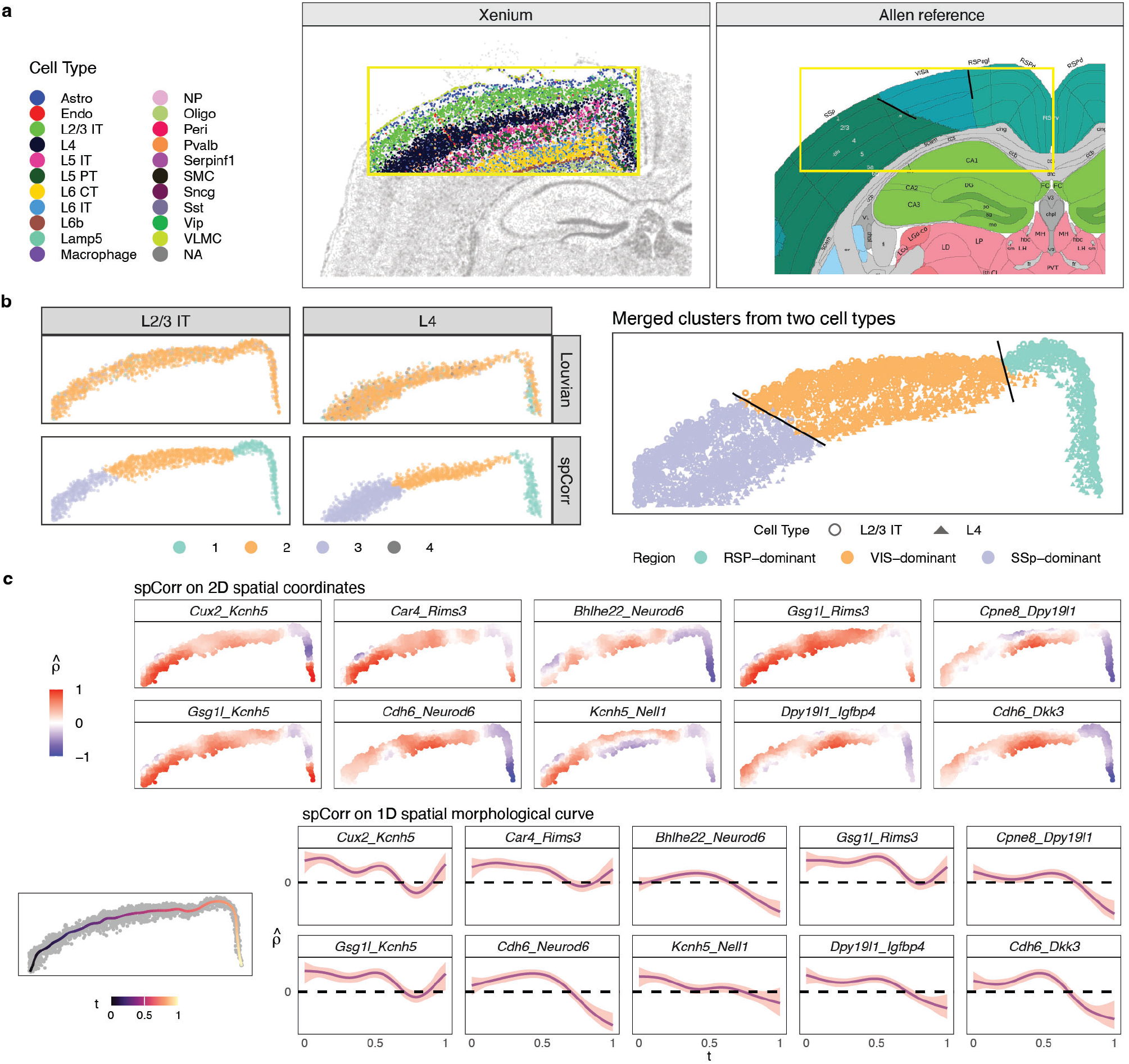
spCorr reveals functional regions within mouse brain cortex layers. **a**. Visualization of mouse brain structure. Left: cell type annotation on a Xenium dataset. The yellow box shows the interested cortex region with clear layer structures. Right: reference structure from Allen Brain Atlas. The yellow box is located approximately in the same place as the one on the left. Solid black lines divide adjacent functional regions across layers. **b**. Comparison of clustering based on gene expression and spatially varying correlation (SVC). Left: clustering results by Seurat Louvain on gene expression and on spCorr-inferred local correlation estimates. While Louvain clusters on gene expression do not show clear patterns, clusters on SVC exhibit three spatially well-separated clusters. Right: alignment of SVC clusters between two adjacent cell-type layers. The three clusters in each cell type are well aligned across layers, and roughly match the three functional domains in the Allen Brain Atlas. **c**. Visualization of spCorr-inferred local correlation estimates. Top: local correlation estimates of the top ten SVC gene pairs in the original two-dimensional space. Bottom: local correlation estimates of the same gene pairs in the one-dimensional spatial curve space. The shadow labels the 95% confidence interval.

To explore computational ways to recover these functional regions, we selected the two most abundant layers, L2/3 IT and L4, for further analysis. We first tried Louvain clustering in Seurat based on the 248 genes; Louvain clustering does not show any patterns (Fig. 5b, top left). This result is not surprising, as the panel genes were chosen to distinguish cell types rather than functional regions. We then applied spCorr to each cell type layer separately. After some gene filtering based on zero proportions to reduce candidate gene pairs, spCorr identified 53 SVC pairs from 8911 gene pairs from L2/3 and 1157 SVC pairs from L4 at the 5% FDR threshold, respectively. Louvain clustering based on local correlation estimates from these significant SVC pairs reveals well-separated clusters horizontal to the cortex structure (Fig. 5b, bottom left). Remarkably, although clustering is performed independently within each cell type layer, the resulting clusters are highly spatially consistent across the two layers and align closely with the three functional regions—RSP, VIS, and SSp (Fig. 5b, right).

**Fig. 5:**
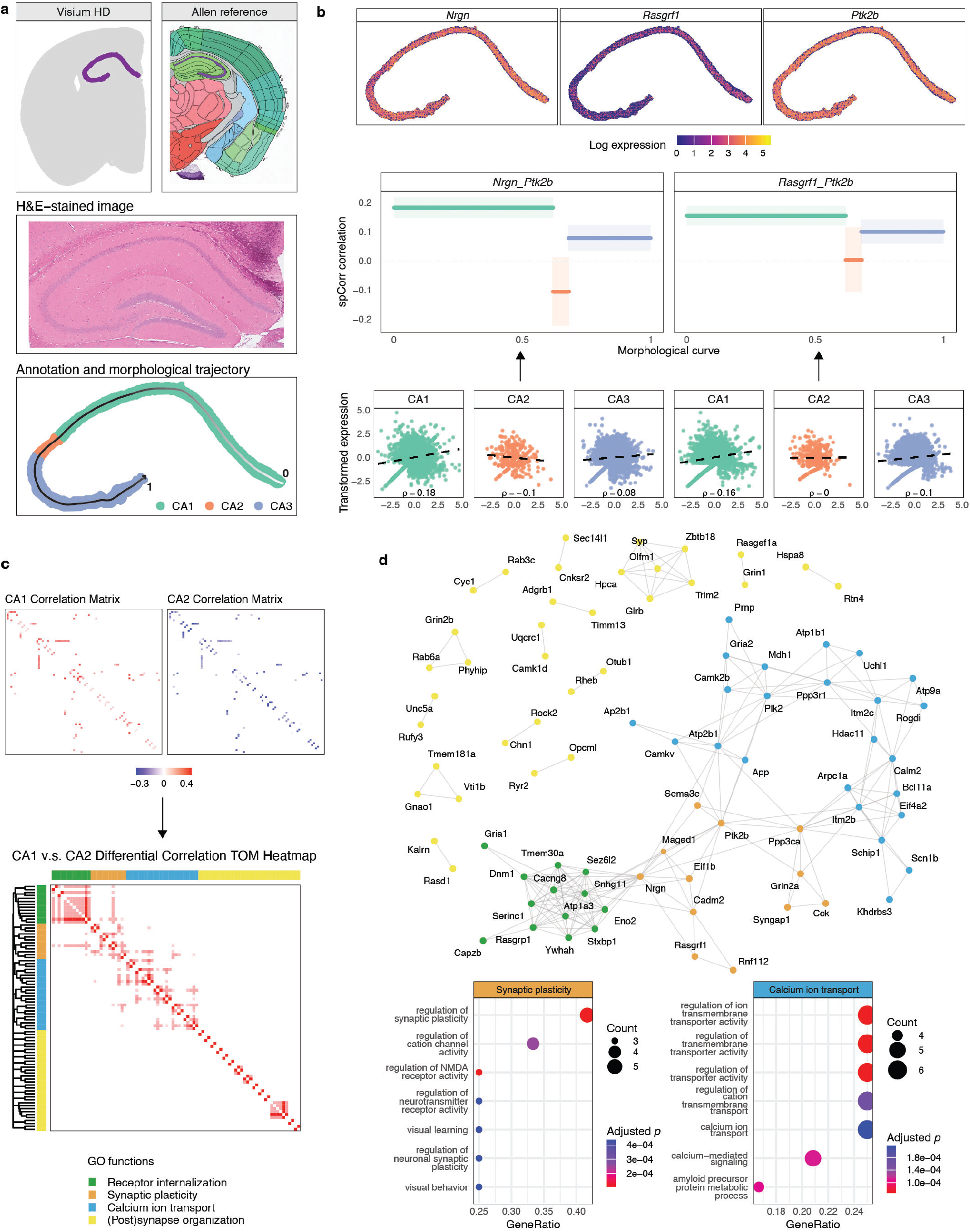
spCorr discovers biologically relevant correlation shift between spatial domains. **a**. Data preprocessing of the Visium HD mouse brain ST dataset. Top: spatial segmentation from BANKSY identifying two adjacent domains corresponding to the pyramidal layer of the hippocampus (aligned with CA1sp-CA3sp regions in the Allen Mouse Brain Atlas). Middle: H&E-stained image of the selected hippocampal region. Bottom: spatial annotation based on CA2 marker gene expression and one-dimensional spatial curve fitted by MorphoGAM along the CA1-CA3 axis. **b**. Local correlation estimation for representative gene pairs. Top: Log-transformed expression levels of *Nrgn, Rasgrf1*, and *Ptk2b*. Middle: spCorr-inferred spot-level correlation estimates for each gene pair, with 95% confidence intervals, modeled using discrete spatial domain annotations and visualized along the spatial curve. Bottom: scatter plot of the transformed gene expressions after the marginal distribution modeling step, showing patterns consistent with the estimated local correlations. **c**. Differential correlation (DC) analysis between CA1 and CA2. Top: Heatmaps of estimated correlation matrices for significantly DC gene pairs, showing broad downregulation in CA2. Bottom: Hierarchical clustering of the differential correlation matrix identified four gene modules enriched for distinct neuronal processes. **d**. Network and functional interpretation of DC results. Top: DC network visualization for gene pairs with statistically significant correlation shifts between CA1 and CA2. Bottom: GO function plot for two representative modules, highlighting synaptic plasticity and calcium ion signaling—biological processes known to differ between CA1 and CA2.

To further visualize the SVC patterns, we plotted the local correlation estimates from 10 top SVC gene pairs in layer L2/3 IT, which have the smallest *p*-values and largest wigglyness (Fig. 5c, top). Notably, the local correlation estimates for most gene pairs exhibit a dramatic shift between the RSP and VIS regions, even when the marginal expression of the individual genes does not display such a pattern (Fig. S6). Considering that the cell-type layer has an elongated shape, we applied ideas of the spatial morphological curve from the

R package MorphoGAM [33], which reduces the spatial coordinate system from 2 dimensions to 1 dimension [33]. spCorr captures similar patterns on this 1D curve as those from 2D space (Fig. 5c, bottom). The 95% confidence intervals provide the local confidence levels of the estimated correlation at each morphological location. While reflecting similar SVC patterns in 2D space, this 1D “reduced” approach dramatically speeds up the computation - 15 times faster than the 2D case for L2-3 IT and 2 times for L4. In summary, spCorr expands the feature space by considering gene interactions when the total gene number is limited. The significant SVC patterns on gene pairs reveal important functional regions that are orthogonal to cell types obtained from regular gene-level analysis.

### 2.5 spCorr discovers biologically relevant correlation shift between mouse hippocampus spatial domains

In the third real-data application, we analyzed a mouse brain hippocampus dataset generated with high-resolution 10x Visium HD technology. The original dataset includes expression measurements for 18,991 genes across 393,543 spots, captured at an aggregated 8 *µ*m bin resolution. We selected spatially distinct tissue structures that correspond to the pyramidal layer of the hippocampus (Fig. 5a, top), yielding a subset of 5,910 spots, as visualized in the corresponding H&E-stained image (Fig. 5a, middle). We further annotated this hippocampal region into CA1, CA2, and CA3 domains based on the expression of canonical marker genes (Fig. S8), aligning with the CA1sp, CA2sp, and CA3sp regions defined in the Allen Mouse Brain reference atlas [34]. Given the slender, curved spatial organization of the CA1-CA3 pyramidal layer, we again applied MorphoGAM [33] to fit a one-dimensional spatial curve that effectively captured the spatial variation along this axis (Fig. 5a, bottom). For gene-level quality control, we retained 964 genes that were expressed in at least 10% of the spots within each of the CA1, CA2, and CA3 domains.

We next applied spCorr to estimate spot-level gene correlations after modeling the marginal distribution given the spatial domain label to adjust average expression differences between the CA1, CA2, and CA3 domains. Specifically, we analyzed 464,166 gene pairs derived from 964 candidate genes, modeling the correlation as a function of the pre-annotated hippocampal domains. Using SVC testing, spCorr identifies 4,873 significant gene pairs at the 5% FDR threshold. Moreover, to illustrate the local correlation estimation, we visualized the log expression of three representative genes -*Nrgn* (Neurogranin), *Rasgrf1* (RAS protein-specific guanine nucleotide-releasing factor 1), and *Ptk2b* (Protein tyrosine kinase 2 beta) (Fig. 5b, top), alongside their spot-level correlation estimates with confidence intervals from spCorr, shown along the spatial morphological curve (Fig. 5b, middle). The domain-specific correlation estimates for each gene pair are consistent with the correlations computed from their transformed expressions, which are obtained after the marginal distribution modeling (step 1) and Gaussian transformation (step 2) in spCorr (Fig. 5b, bottom).

To investigate how active gene modules and pathways differ between hippocampal domains, we extended our analysis to differential correlation (DC) testing. A longstanding question in hippocampal neurobiology is whether CA2 represents a functionally distinct subfield or merely serves as a transition zone between CA1 and CA3. Although CA2 is anatomically positioned between CA1 and CA3 and shares structural features with both, accumulating evidence from pathological and molecular studies supports its classification as a distinct region [35, 36]. Motivated by this, we used spCorr to perform DC testing to determine whether gene correlation patterns differ significantly between CA1 and CA2.

The DC test between CA1 and CA2 identifies 69 significant gene pairs at the 5% FDR threshold, involving 83 unique genes. A heatmap of the estimated correlation matrices for these gene pairs in CA1 and CA2 reveals a clear down-regulation in correlation from CA1 to CA2 (Fig. 5c, top). To systematically quantify these differences, we computed a differential correlation matrix by subtracting the estimated correlations in CA2 from those in CA1. We then applied weighted gene correlation network analysis (WGCNA) to the differential correlation matrix, identifying four gene modules via hierarchical clustering of the topological overlap matrix (TOM) [8]. To investigate the biological functions associated with these modules, we carried out Gene Ontology (GO) enrichment analysis for each cluster. The results reveal that the modules are significantly enriched for GO terms associated with key neuronal processes, including receptor internalization, synaptic plasticity, calcium ion transport, and the organization of synaptic and postsynaptic structures (Fig. 5c, bottom). To provide a network-based perspective on these gene correlation changes, we further proposed the gene differential correlation network, where edges represent gene pairs with statistically significant differential correlation, and edge weights reflect the magnitude of correlation difference between CA1 and CA2 (Fig. 5d, top). Previous studies have shown that CA2 neurons exhibit markedly reduced synaptic plasticity and distinct calcium signaling dynamics relative to CA1 [37–39]. These observations are consistent with our GO enrichment results, as two of the identified modules are significantly associated with synaptic plasticity and calcium ion transport (Fig. 5d, bottom), reconfirming the biological distinctiveness of the CA2 region.

Additionally, to demonstrate the necessity of the marginal distribution modeling detp (step 1)in the spCorr pipeline, where marginal distributions were modeled conditional on spatial domain annotations (CA1, CA2, CA3), we conducted a comparative analysis in which this adjustment was omitted. Specifically, we repeated the DC analysis between CA1 and CA2 without accounting for average expression differences between domains in the marginal distribution modeling step. In this case, the analysis still identifies four distinct gene modules (Fig. S9a, b); however, the functional enrichment results are less aligned with the known biological distinctions between the CA1 and CA2 regions (Fig. S9c). These findings underscore the importance of properly modeling marginal distributions to reveal biologically meaningful spatially varying correlation patterns.

Although our primary analyses have been focused on the DC analysis between spatial domains in ST data, the spCorr framework is equally applicable to DC analysis between cell types in scRNA-seq data. To demonstrate this generalizability, we applied spCorr to a mouse hippocampus scRNA-seq dataset [40]. Focusing on annotated pyramidal neurons (CA1, CA2, and CA3 principal cells), we observed similar DC patterns between CA1 and CA2. These DC gene pairs are again enriched for functional categories related to synaptic plasticity and calcium ion transport (Fig. S10), similar to the findings from ST data. This independent analysis reinforces the biological relevance and cross-platform robustness of the DC signals identified by spCorr.

## 3 Discussion

We propose a novel statistical framework, spCorr, designed to detect spatially varying correlation (SVC) in spatial transcriptomics data. Unlike existing methods that primarily rely on non-parametric local smoothing, spCorr emphasizes rigorous and interpretable inference through a flexible parametric model. A key innovation of spCorr is its transformation of the pairwise correlation estimation problem into a univariate regression problem. Specifically, spCorr first models each gene’s marginal distribution conditional on potential confounding covariates and transforms gene expression values into standard Gaussian variables. Leveraging Gaussian copula theory, it then constructs a gene-pair cross-product variable and applies a quasi-GAM regression with spatial coordinates as predictors. Finally, spCorr computes *p*-values in SVC testing using asymptotic theory for quasi-GAMs, and derives the local correlations as an explicit function of spatial location.

Extensive simulations demonstrate that spCorr achieves superior false discovery rate (FDR) control and statistical power compared to state-of-the-art methods like scHOT, SpatialDM, and SpatialCorr. Notably, spCorr is the only method tested that consistently yields well-calibrated *p*-values and effectively controls the FDR, even when the data do not perfectly match its model assumptions. This statistical robustness, combined with the flexibility of the GAM framework, enables spCorr to attain the highest power and AUROC across all benchmarks. In real-data applications, spCorr uncovered spatially heterogeneous regulatory patterns and identified spatial domains that were not detectable by conventional gene-level analyses.

A central assumption of spCorr is the Gaussian copula model, which posits that the correlation structure among genes can be represented via a multivariate Gaussian distribution. Our prior work, scDesign3 [25], established that this model provides a robust and accurate approximation of high-dimensional count data. A unique advantage of the copula framework is its ability to decouple marginal distributions from the dependence structure, allowing spCorr to accommodate a wide range of distributional forms, including Negative Binomial (default for scRNA-seq), Poisson, Zero-Inflated Negative Binomial, Zero-Inflated Poisson, and Gaussian. This decoupling also facilitates the adjustment for arbitrary confounders by modeling each gene’s marginal distribution conditional on covariates-akin to *partial correlation* in the linear setting.

For instance, in our real-data analysis comparing the CA1 and CA2 domains of the hippocampus, we first regressed out the domain labels to equalize each gene’s marginal distribution across regions. This step prevents correlation estimates from being confounded by differences in mean expression levels. Omitting this adjustment would lead the correlation estimates to reflect average expression shifts rather than true domain-specific coordination, resulting in the identification of gene modules enriched in generic biological processes (e.g., ATP metabolism and mitochondrial organization; see Fig. S9). This issue, a form of *Simpson’s paradox*, underscores the critical importance of adjusting for confounding effects. While spCorr provides flexibility for marginal modeling, we strongly recommend that users carefully evaluate covariates based on their biological question. In extreme cases, one can even model the marginal distribution as a function of spatial coordinates to remove all spatial effects on gene expression means. A recent preprint adopted this approach to isolate cell-intrinsic coordination by regressing out spatial effects before computing correlation [41].

There are several important future directions. Currently, spCorr focuses on gene correlations from spatial transcriptomics. However, to construct more comprehensive gene regulatory networks, integration with additional omics layers—such as chromatin accessibility from spatial ATAC-seq—is necessary. For example, peak-gene correlations have been used to infer cis-regulatory interactions [42, 43]. Extending spCorr to quantify peak-gene correlations in spatial multi-omics data [20, 44] is a natural and feasible next step, thanks to the copula model’s flexibility in handling heterogeneous marginal distributions.

Second, while spCorr uses regression splines (e.g., thin plate splines and cubic splines) to capture spatial variation, this assumes smooth spatial surfaces. In reality, tissue structure often leads to abrupt transitions or boundaries in expression patterns. To accommodate such features, future versions of spCorr could incorporate histological images (e.g., Hematoxylin & Eosin staining) to inform spatial discontinuities. In summary, extending spCorr to integrate multi-modal spatial data holds great promise for advancing our understanding of spatial gene regulation.

## 4 Method

### 4.1 Mathematical notations

We define a *spot* as a single observation in a spatial transcriptomics dataset. Depending on the technology, a spot may correspond to a capture area (e.g., 10x Visium), a segmented cell (e.g., 10x Xenium), or a bin (e.g., 10x Visium HD). spCorr requires three input matrices: a spot-by-gene expression matrix **Y**, a spatial coordinate matrix **S**, and an optional spot-by-covariates matrix **X** (Fig. 1, left).

First, let **Y** = [*Y*_*ij*_] ∈ ℝ^*n×m*^ denote the gene expression matrix, where *n* is the number of spots and *m* is the number of genes. While in this manuscript, we mainly discuss the gene count matrix **Y** = [*Y*_*ij*_] ∈ ℕ^*n×m*^, the method is also generalizable to continuous data such as log-transformed gene expression. Second, let **S** = [***s***_1_, …, ***s***_*n*_]^⊤^ ∈ ℝ^*n×*2^ denote the spatial coordinate matrix, where each ***s***_*i*_ = (*s*_*i*1_, *s*_*i*2_)^⊤^ specifies the two-dimensional spatial coordinate of spot *i*. Third, let **X** = [***x***_1_, …, ***x***_*n*_]^⊤^ ∈ ℝ^*n×q*^ denote the matrix of user-defined covariates, where each ***x*** _*i*_ = (*x*_*i*1_, …, *x* _*iq*_)^⊤^ captures *q* covariates for spot *i* that are associated with gene expression (e.g., library size, spot annotation) that may confound the correlation estimation based on the scientific questions.

### 4.2 Gaussian copula model

The spCorr model is formulated using a Gaussian copula (Fig. 1, middle):

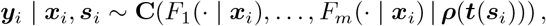

where *F*_*j*_ (· | ***x***_*i*_) is the conditional marginal distribution of gene *j* given ***x***_*i*_, and **C** is the Cumulative Distribution Function (CDF) of a multivariate standard Gaussian with correlation matrix ***ρ***(***t***(***s***_*i*_)).

The key feature of spCorr is that the correlation matrix is not fixed but varies with spatial predictors ***t***(***s***_*i*_), which may represent two-dimensional spatial coordinates, one-dimensional spatial curves, or discrete spatial domains. For a gene pair (*j, k*), the local correlation at spot *i* is modeled as

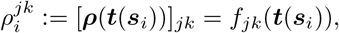

where *f*_*jk*_ describes how local copula correlation between gene *j* and gene *k* varies as a function of spatial structure. This pairwise approach enables interpretable and efficient inference of spatial gene-gene relationships.

### 4.3 The spCorr algorithm

The Gaussian copula model defines the overall framework of spCorr. In the following, we describe the implementation of the spCorr algorithm (Fig. 1, middle).

#### Marginal distribution modeling

For each gene *j* = 1, …, *m* at every spot *i* = 1, …, *n*, spCorr uses a Generalized Linear Model (GLM) to model the distribution of the observed expression value *Y*_*ij*_ conditional on spot-level covariate ***x***_*i*_:

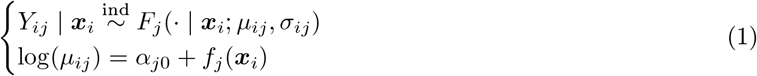

where *F*_*j*_(·; *µ*_*ij*_, *σ*_*ij*_) denotes the cumulative distribution function (CDF) of the assumed conditional distribution of *Y*_*ij*_ with mean parameter *µ*_*ij*_ and scale parameter *σ*_*ij*_. The function *f*_*j*_(***x***_*i*_) captures the marginal effect of the spot-level covariates ***x***_*i*_. The specific form of *F*_*j*_ and *f*_*j*_(·) depends on the data distribution and the format of covariates, respectively. For instance, if ***Y*** is a count matrix, and ***x***_*i*_ encodes categorical spatial domain annotations, then *F*_*j*_ is the CDF of a Negative Binomial (NB) distribution, and 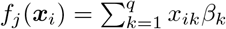 is a regression formula on spatial domain annotation. The choice *f*_*j*_ is essentially flexible and may take a nonlinear format, such as a spline, which will change the GLM into a Generalized Additive Model (GAM). spCorr fits the GLM using the R package `mgcv`, which estimates model parameters by penalized-restricted maximum likelihood estimation (MLE). The fitted CDF for gene *j* is denoted by 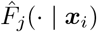.

#### Gaussian transformation

To enable Gaussian copula-based correlation estimation, the observed expression values *Y*_*ij*_ are transformed to continuous standard Gaussian variables. Because ST data is often discrete (counts), we first apply a randomized quantile transformation to obtain a continuous uniform variable [25, 45]:

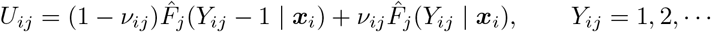

where *ν*_*ij*_ *∼* Uniform[0, 1] is sampled independently. Each *U*_*ij*_ is then transformed into a standard Gaussian variable using the inverse CDF of the standard Gaussian distribution:

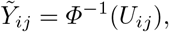

where *Φ*(·) is the CDF of the standard Gaussian distribution. The resulting 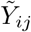 approximately follows a standard Gaussian distribution.

#### Local correlation estimation

For any gene pair (*j, k*) at spot *i*, we consider the bivariate distribution of the transformed expressions 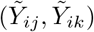, where both variables approximately follow standard Gaussian distributions: 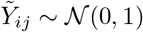 and 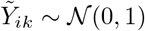. Their dependence is characterized by the Pearson correlation coefficient:

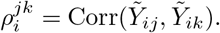

Based on the theory of the Gaussian copula [46], this quantity is directly related to Kendall’s *τ* of the original variables via:

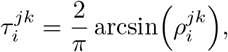

implying that modeling 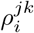 spatially enables estimation of the local Kendall’s *τ* across the space. To this end, we define a new random variable 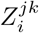, the cross-product of transformed expressions:

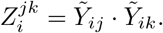

which, under bivariate normality, satisfies:

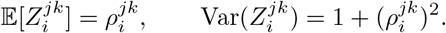

This new random variable, 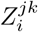, allows us to convert the modeling of a bivariate distribution into a univariate distribution. spCorr models the spatial variation in 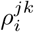 as a function of a spatial predictor ***t***(***s***_*i*_):

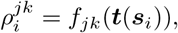

where ***t***(***s***_*i*_) is derived from the spatial location ***s***_*i*_. Depending on the application context, this spatial predictor may take one of the following forms:

– a two-dimensional spatial coordinate: ***t***(***s***_*i*_) = (*s*_*i*1_, *s*_*i*2_) ∈ ℝ^2^,
– a one-dimensional spatial morphological curve: ***t***(***s***_*i*_) = *t*_*i*_ ∈ ℝ,
– or a discrete spatial domain label: ***t***(***s***_*i*_) ∈ {0, 1, …}.

While it is possible to use MLE to estimate *f*_*jk*_, the distribution of *Z*_*ij*_ is very complicated, making the computation challenging. However, the variance of *Z*_*ij*_ has a simple relation to its mean, which reminds us of the quasi-likelihood theory as an alternative inference strategy [47]. To estimate 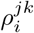, we model the conditional distribution of 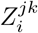 given ***t***(***s***_*i*_) using a quasi-likelihood generalized additive model (Quasi-GAM). Based on the known mean and variance structure of 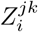, the corresponding quasi-likelihood function is:

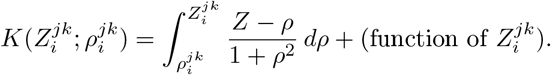

We then fit a Quasi-GAM by maximizing the quasi-likelihood for each gene pair (*j, k*) as:

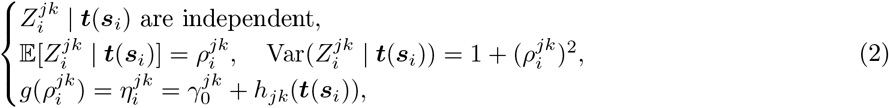

where *g*(·) = artanh(·) is the hyperbolic arctangent link function mapping (−1, 1) to ℝ, and 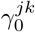 is the intercept specific to the gene pair (*j, k*).

The spatial smoother *h*_*jk*_(***t***(***s***_*i*_)) is defined flexibly depending on the type of spatial predictor ***t***(***s***_*i*_), which allows spCorr to adapt to diverse spatial structures:

– **Spatial coordinates** (***t***(***s***_*i*_) ∈ ℝ^2^): When spatial location is represented by two-dimensional coordinates, *h*^*jk*^(·) is modeled using a thin-plate spline. Specifically, it is expressed as a linear combination of radial basis functions: 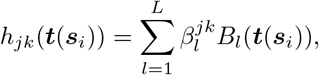

where *B*_*l*_: ℝ^2^ → ℝ are pre-defined radial basis functions, and 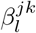 denotes the coefficient for the *l*-th basis function corresponding to gene pair (*j, k*).
– **Spatial curves** (***t***(***s***_*i*_) ∈ ℝ): When spatial variation is captured by a one-dimensional spatial curve, *h*^*jk*^(·) is modeled using a univariate cubic spline:

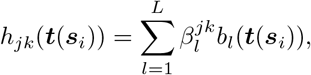

where *b*_*l*_: ℝ → ℝ are cubic spline basis functions, and 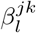 are the corresponding coefficients for gene pair (*j, k*). Such one-dimensional spatial curves may be defined manually (based on tissue morphology) or inferred computationally (e.g., MorphoGAM [33]) and then used as the spatial predictors.
– **Spatial domains** (***t***(***s***_*i*_) ∈ {0, 1, …, *D* − 1}): When space is partitioned into discrete domains, *h*^*jk*^(·) is modeled as a domain-specific effect using a one-hot encoding: 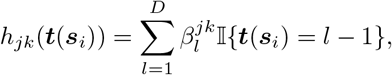

where 𝕀{·} is the indicator function, and 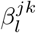 represents the corresponding coefficients in domain *l* for gene pair (*j, k*).

Model fitting is performed using the `mgcv`package in `R`. For continuous spatial predictors (e.g., spatial coordinates or morphological curves), the dimension of the basis, denoted by *L*, controls the flexibility of the smoother. The exact choice of *L* is typically not critical as long as it is sufficiently large, since spCorr employs penalized maximum likelihood estimation by default to regularize the smoother. This regularization imposes smoothness over space, allowing the model to naturally borrow information from neighboring spatial locations. Disabling the penalty can simplify inference by making *p*-value computation more straightforward, as it avoids uncertainty due to smoothing parameter estimation. However, doing so may reduce model flexibility and increase the risk of overfitting. For discrete spatial predictors (e.g., spatial domain labels), no penalization is applied, as the smoother reduces to a set of categorical effects.

The final correlation estimate for gene pair (*j, k*) at spot *i* is given by

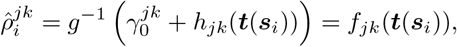

providing an interpretable function of spatial location for the local correlation between genes *j* and *k*, as the model explicitly links correlation strength to spatial predictors such as spatial coordinates, spatial curves, or domain labels.

**Hypothesis testing** spCorr supports two types of statistical testing to assess spatial correlation variation for each gene pair (*j, k*):

– **Spatially varying correlation (SVC)** testing, which evaluates whether correlation varies over continuous spatial predictors (e.g., spatial coordinates or curves).
– **Differential correlation (DC)** testing, which compares the correlation between discrete spatial predictors (e.g., spatial domain labels).

##### SVC testing

Let 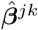 ∈ℝ^*K*^ denote the estimated coefficients of the smooth function *h*_*jk*_ from the Quasi-GAM model. Under regularity conditions, these coefficients satisfy asymptotic normality

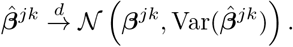

The null and alternative hypotheses in testing the SVC are

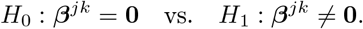

A significant result indicates that the local correlation 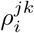 exhibits spatial heterogeneity.

##### DC testing

Let *d*_*i*_ ∈ {0, 1, …} denote the discrete spatial domain label for spot *i*, and let *d*^1^ and *d*^2^ be two specific domains under comparison. In the Quasi-GAM framework, DC testing evaluates the difference in estimated domain-specific coefficients:

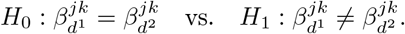

Let 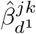 and 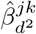 denote the estimated coefficients for domains *d*^1^ and *d*^2^, respectively. The Wald test statistic is:

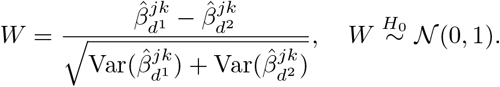

The resulting two-sided *p*-value determines significance; a small *p*-value indicates evidence of differential correlation between the two spatial domains.

##### FDR control

For both SVC and DC testing, spCorr applies the Benjamini–Hochberg (BH) procedure [48] to control the FDR across all tested gene pairs. By default, the FDR threshold is set to 0.05.

**Confidence interval** spCorr provides spot-level correlation estimates 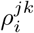 with uncertainty quantification, i.e., confidence intervals for each 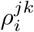. Given the fitted value 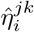 and its estimated standard error 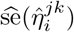 from the quasi-GAM model (Eq. 2), we first construct a (1 − *α*) confidence interval for the linear predictor as

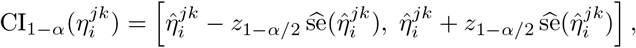

where *z*_1−*α/*2_ is the (1 − *α/*2) quantile of the standard normal distribution.

Then, applying the inverse link function *g*^−1^(·) = tanh(·) maps the interval back to the correlation scale, yielding the confidence interval for 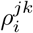 as

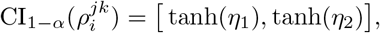

where 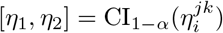.

### 4.4 Downstream analysis

The output of the spCorr algorithm enables several types of downstream analysis (Fig. 1, right).

**SVC/DC identification** spCorr identifies informative gene pairs that show significant correlation changes across continuous space (SVC) or between discrete domains (DC). For these gene pairs, the estimated spot-level correlation can be visualized to inspect how functionally related genes correlate across spatial contexts.

**Spatial clustering** The estimated local correlation matrices (spot by gene pair) from spCorr can be used for clustering analyses. For example, Louvain clustering applied to these matrices can reveal spatial heterogeneity that may not be apparent from gene expression alone.

**Gene correlation network analysis** spCorr also supports network-level analysis. For a given correlation matrix (e.g., from differential correlation), we apply Weighted Gene Correlation Network Analysis (WGCNA) [8]. The correlation matrix is first transformed into an adjacency matrix, followed by computation of the topological overlap matrix (TOM) and corresponding dissimilarity. Hierarchical clustering is then applied to the dissimilarity matrix, and gene modules are identified by dynamic tree cutting.

## 5 Data Availability

The human dorsolateral prefrontal cortex (DLPFC) 10x Genomics Visium dataset [27] used for simulations is accessible through the R package `spatialLIBD`[49]. The human oral squamous cell carcinoma 10x Visium dataset (*sample 2* from [21]) can be downloaded from https://doi.org/10.6084/m9.figshare.20304456. v1. The mouse brain cortex 10x Xenium dataset is provided by 10x Genomics as part of the “Fresh Frozen Mouse Brain for Xenium Explorer Demo,” available at https://www.10xgenomics.com/datasets/fresh-frozen-mouse-brain-for-xenium-explorer-demo-1-standard. The mouse hippocampus 10x Visium HD dataset is also provided by 10x Genomics as part of the “Visium HD Spatial Gene Expression Library, Mouse Brain (FFPE),” available at https://www.10xgenomics.com/datasets/visium-hd-cytassist-gene-expression-libraries-of-mouse-brain-he. The single-cell RNA-seq hippocampus dataset from [40] is available as a processed Seurat object at https://www.dropbox.com/scl/fi/trw3ujacyi33tawjgd5cr/mouse_hippocampus_reference.rds?rlkey=3gns05ok7qc9cu48voq0atiyp&e=1&dl=0, with raw count matrices downloadable from the DropViz website (http://dropviz.org/). The TRRUST v2 database [28] used to select transcription factor–target gene pairs can be accessed at https://www.grnpedia.org/trrust/. A detailed summary of all datasets and preprocessing steps is provided in Supplementary Methods S1.2.

## 6 Code Availability

The R package spCorr is available at https://github.com/chexjiang/spCorr.

## Supplementary Material for

## S1 Supplementary Methods

### S1.1 Simulation designs

#### Simulation setting 1 based on semi-synthetic ST data

To benchmark SVC identification methods in terms of FDR control, statistical power, and computational efficiency, we required ground truths for both SVC and non-SVC gene pairs. To this end, we adopted a semi-synthetic simulation framework that integrates real ST data with user-defined spatial correlation structures, providing a balance between biological realism and controlled evaluation (Fig. S1).

Our simulations were based on ST data from the human dorsolateral prefrontal cortex (DLPFC), specifically sample 151676 generated by 10x Genomics Visium, which contains expression profiles of 33,538 genes across 3,611 spots [27]. The spots span six cortical layers and white matter (WM), with the following distribution: L1 (273 spots), L2 (253), L3 (989), L4 (218), L5 (673), L6 (692), and WM (513) (Fig. S2).

This real dataset provided realistic library sizes, gene-level means, and variances. Because the true spatial correlation structure between gene pairs is unknown in real data, we introduce controlled latent correlation patterns to define known SVC and non-SVC gene pairs.

We selected the top 400 spatially variable genes based on Moran’s I statistic and randomly constructed 200 gene pairs for simulation. The expression data for each gene pair were generated using a Poisson-lognormal model, which enabled flexible specification of spatially varying correlation structures while preserving realistic UMI count distributions. The simulation process involved three key steps:

– **Library size estimation:** For each spot, the library size *ℓ*_*i*_ was computed from the real data.
– **Latent parameter estimation:** For each gene, the region-specific mean *µ*_*j*_(*d*_*i*_) of the log-normal distribution was estimated from the real data, using manual annotations of cortical layers as spatial domains *d*_*i*_.
– **Spatial correlation modeling:** For each gene pair (*j, k*), latent Gaussian variables (*λ*_*ij*_, *λ*_*ik*_) were sampled from a bivariate normal distribution with a user-specified correlation function *ρ*(***s***_*i*_), which was either spatially varying (for SVC pairs) or constant (for non-SVC pairs).

Formally, for each spot *i*, the latent variables were drawn as:

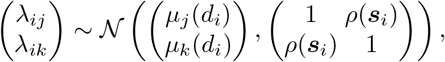

and the observed UMI counts were generated via:

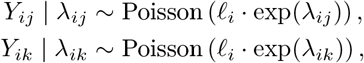

where ***s***_*i*_ = (*s*_*i*1_, *s*_*i*2_) represents the spatial coordinates of spot *i*.

Under this model, the expressions of SVC gene pairs were simulated by allowing *ρ*(***s***_*i*_) to vary smoothly across spatial coordinates ***s***_*i*_, while the expressions of non-SVC gene pairs were simulated with a constant *ρ* across all spots. The resulting Poisson-distributed counts captured realistic gene-level variability and spatial structure observed in ST data.

For each simulation setting, we generated 30 independent replicates. The resulting simulated datasets provided known SVC and non-SVC gene pairs for rigorous and reproducible evaluation of SVC identification methods.

##### Fixed spot number setting

The simulated dataset described above, containing 3,611 spots across L1-L6 and WM, was used to evaluate *p*-value calibration, FDR control, statistical power, and AUROC, as shown in Fig. 2a-d.

##### Varying spot number setting

To evaluate computational scalability (Fig. 2e), we varied the number of spots by selecting subsets of increasing size:

– WM + L6 (1,205 spots),
– WM + L6 + L5 (1,878 spots),
– WM + L6 + L5 + L4 + L3 (3,085 spots),
– WM + L6 + L5 + L4 + L3 + L2 + L1 (3,611 spots).

#### Compared methods and evaluation metrics

In the benchmarking semi-synthetic ST dataset, we compared the performance of spCorr (both penalized and unpenalized GAM regression, using thin-plate spatial spline with *K* = 50) against three existing SVC identification methods: scHOT (version 0.99.7, with default local weighting shape and span), SpatialDM (version 0.1.0, using scale factor *l* = 0.2 and cut-off *co* = 0.2), and SpatialCorr (version 1.2.0, using within-region test with kernel bandwidth *γ* = 100).

We evaluated five performance metrics across methods, corresponding to different panels of Fig. 2:

(1) Calibration of *p*-values under the null (non-SVC gene pairs), evaluated by quantile-quantile (Q-Q) plots against the expected Uniform[0,1] distribution, and histograms of observed *p*-values (Fig. 2a). Well-calibrated *p*-values are expected to follow the Uniform[0,1] distribution.
(2) FDR control, assessed by the actual FDR achieved at a target FDR threshold of 0.05 using the Benjamini-Hochberg procedure (*p* ≤ 0.05 after adjustment) (Fig. 2b). Accurate FDR control requires the actual FDR to remain close to the target level.
(3) Statistical power, measured as the proportion of true SVC gene pairs detected under the actual FDR = 0.05 cutoff (Fig. 2c). Higher power indicates greater sensitivity for detecting SVC.
(4) Discriminative ability, quantified by receiver operating characteristic (ROC) curves and the corresponding area under the ROC curve (AUROC), which reflects the ability to distinguish SVC from non-SVC gene pairs (Fig. 2d).
(5) Computational efficiency, benchmarked by the log-transformed runtime (seconds per gene pair) as a function of the number of spots (Fig. 2e). Better scalability is indicated by lower runtime as the dataset size increases.

For SpatialCorr, we additionally investigated the sensitivity of performance with respect to the band-width parameter *γ*. In SpatialCorr, *γ* is the bandwidth in the Gaussian kernel function

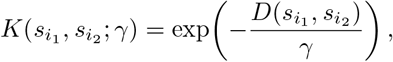

where 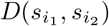 is the Euclidean distance between spots *i*_1_ and *i*_2_. Because *γ* is compared directly to the distances in the original coordinate scale, its choice can substantially influence results. In our simulation settings, the spatial coordinates ranged over *s*_*i*1_ ∈ (3096, 11062) and *s*_*i*2_ ∈ (2437, 11195). Under this scale, the default *γ* = 5 in the SpatialCorr Python package leads to numerical errors: too small *γ* drives kernel values near zero, resulting in a nearly singular covariance matrix. To avoid this, we tested *γ* 1∈ {0^2^, 10^3^, 10^4^, 10^5^, 10^6^}, which are more comparable to the coordinate scale, and evaluated results accordingly. Our analysis indicates that varying *γ* across this range has little impact on FDR, power, or AUROC (Fig. S4).

### S1.2 Real data analysis

#### Collection of real data

We used three ST datasets generated by different technologies: an OSCC tumor Visium dataset, a mouse brain cortex Xenium dataset, and a mouse brain hippocampus Visium HD dataset. The OSCC tumor Visium dataset (*sample 2* from the study [21]) was downloaded from https://doi.org/10.6084/m9.figshare.20304456.v1. The mouse brain cortex Xenium dataset was provided by 10x Genomics as part of the “Fresh Frozen Mouse Brain for Xenium Explorer Demo,” available at https://www.10xgenomics.com/datasets/fresh-frozen-mouse-brain-for-xenium-explorerdemo-1-standard. The mouse brain hippocampus Visium HD dataset was provided by 10x Genomics as part of the “Visium HD Spatial Gene Expression Library, Mouse Brain (FFPE),” and downloaded from https://www.10xgenomics.com/datasets/visiumhd-cytassistgene-expression-libraries-of-mouse-brain-he. We also used one scRNA-seq hippocampus dataset produced by [40], available for download as a processed Seurat object at https://www.dropbox.com/scl/fi/trw3ujacyi33tawjgd5cr/mouse_hippocampus_reference.rds?rlkey=3gns05ok7qc9cu48voq0atiyp&e=1&dl=0, with the raw count matrices available from the DropViz website at http://dropviz.org/. Additionally, we used the TRRUST v2 database [28] to select TF-target gene pairs, available at https://www.grnpedia.org/trrust/.

#### Data preprocessing

##### OSCC tumor Visium data

This dataset consists of ST data from HPV-negative oral squamous cell carcinoma (OSCC) sequenced using the 10x Genomics Visium platform. We used sample 2 as the main example, which contains expression measurements for 15,624 genes across 1,749 spots with pathologist annotations (Fig. 3a), and was originally normalized using `sctransform`[50]. For spot-level selection, we excluded two outlier spots that lacked pathologist annotations. For gene-level filtering, we restricted the analysis to TF-target pairs from the TRRUST v2 database. We further refined this set by removing those TF-target pairs classified as “Repression” and filtered out genes with expression in fewer than 10% of spots. This resulted in 1,316 candidate TF-target gene pairs across 1,747 spots.

##### Mouse brain cortex Xenium data

This dataset consists of ST data from the mouse brain measured by 10x Genomics Xenium technology, including expression measurements for 248 genes across 36,553 cells by pre-defined cell segmentations. For spot-level selection, we followed the Seurat tutorial https://satijalab.org/seurat/articles/seurat5_spatial_vignette_2. We first selected a sub-region that mainly contained the cortex region. We then used two cell-type annotation method, RCTD [51] and SingleR [52], to obtain a set of highly reliable cell-type labels. We further focused on the annotated L2/3 IT and L4 layers. Given the slender structures of the cortex layer, we applied MorphoGAM (with default parameters) [33] to fit a one-dimensional spatial curve that effectively captured the spatial variation along this axis (Fig. 4c, bottom). For gene-level quality control, we retained genes that were expressed in at least 20% of the spots within each of the L2/3 IT and L4 layers.

##### Mouse brain hippocampus Visium HD data

This dataset consists of ST data from the mouse brain sequenced using the 10x Genomics Visium HD technology, including expression measurements for 18,991 genes across 393,543 spots, captured at 8 *µ*m resolution. For spot-level selection, we first applied the BANKSY algorithm (`lambda = 0.8, k_geom = 50`) [53] to segment the tissue into 31 spatial domains (Fig. S7). From these, we selected two adjacent domains (25 and 27) corresponding to the pyramidal layer of the hippocampus (Fig. 5a, top), yielding a subset of 5,910 spots, as visualized in the corresponding H&E-stained image (Fig. 5a, middle). We further annotated this hippocampal region into CA1, CA2, and CA3 domains based on the expression of canonical marker genes (Fig. S8), aligning with the CA1sp, CA2sp, and CA3sp regions defined in the Allen Mouse Brain reference atlas [34]. Given the curved spatial organization of the CA1-CA3 pyramidal layer, we applied MorphoGAM (with default parameters) [33] to fit a one-dimensional spatial curve that effectively captured the spatial variation along this axis (Fig. 5a, bottom). For gene-level quality control, we retained 964 genes that were expressed in at least 10% of the spots within each of the CA1, CA2, and CA3 subfields.

#### Gene Ontology enrichment analysis

We performed Gene Ontology (GO) enrichment analysis for biological processes using the R package clusterProfiler (version 4.6.2). The `enrichGO`function was used with the ontology set to `“BP”`, and gene sets with a Benjamini-Hochberg-adjusted *p*-value of ≤ 0.05 were considered significant.

#### Weighted gene correlation network analysis

We performed weighted gene correlation network analysis (WGCNA) [8] on the differential correlation (DC) matrix using the R package WGCNA (version 1.73). The DC matrix was first converted to an adjacency matrix, and the topological overlap matrix (TOM) and corresponding dissimilarity were computed using the `TOMsimilarity`function. Hierarchical clustering was then applied to the dissimilarity matrix using the `hclust`function, and gene modules were identified by dynamic tree cutting using the `cutreeDynamic`function.

#### Heatmaps

We generated correlation heatmaps with genes ordered by hierarchical clustering. The heatmaps were created using the `pheatmap`function from the R package pheatmap (version 1.0.13).

## S2 Supplementary Tables and Figures

**Table S1:** Comparison of SVC identification methods based on key properties.

| Method | Interpretability | Covariate adjust | Count modeling | Theoretical $p$ |
| --- | --- | --- | --- | --- |
| SpatialDM [22] | × | × | × | ✓✗ |
| scHOT [23] | × | × | × | × |
| SpatialCorr [24] | × | ✓ | × | × |
| **spCorr** | ✓ | ✓ | ✓ | ✓ |

Property 1: Interpretable modeling of the correlation as an explicit function of spatial predictors Property 2: Covariate adjustment; if confounding effects can be adjusted by the model

Property 3: Direct modeling of count data without log transformation

Property 4: Theoretical *p*-values. The *p*-value calculation is based on the asymptotic distribution and thus does not require a permutation test.

For each method and each property, a checkmark, half-checkmark, or cross means that the method satisfies, partially satisfies, or does not satisfy the property, respectively.

**Fig. S1:**
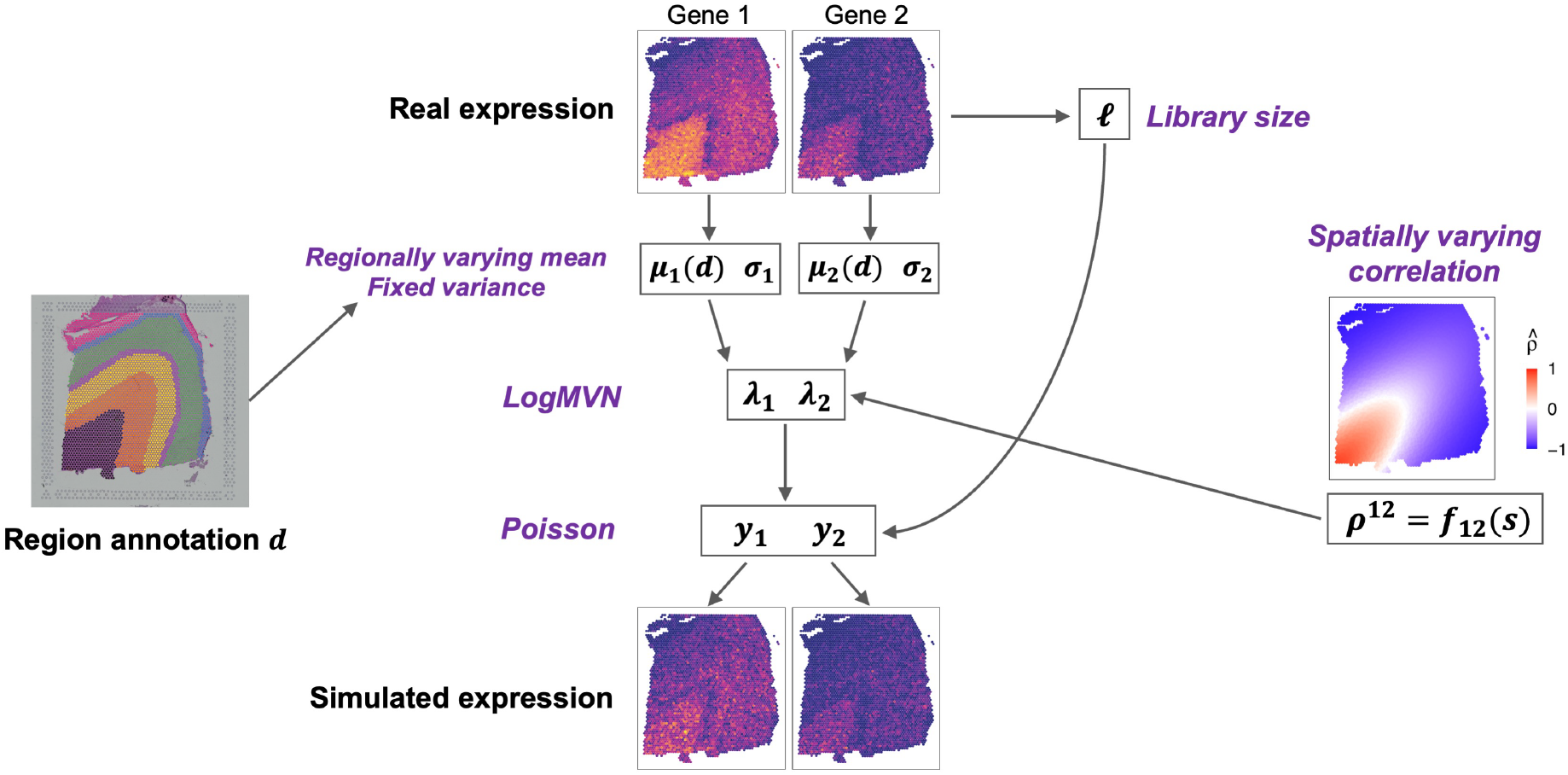
Simulation setting 1: Overview of the semi-synthetic simulation framework. To balance biological realism with controlled ground truth, simulated ST data were generated by combining real Visium data from the human dorsolateral prefrontal cortex with user-defined spatial correlation structures. A Poisson-lognormal model was used, where gene-specific means, variances, and library sizes were estimated from real data, while spatially varying or constant gene correlations were introduced at the latent level to define SVC or non-SVC gene pairs. This hybrid design enabled rigorous evaluation of spCorr’s performance under known correlation patterns.

**Fig. S2:**
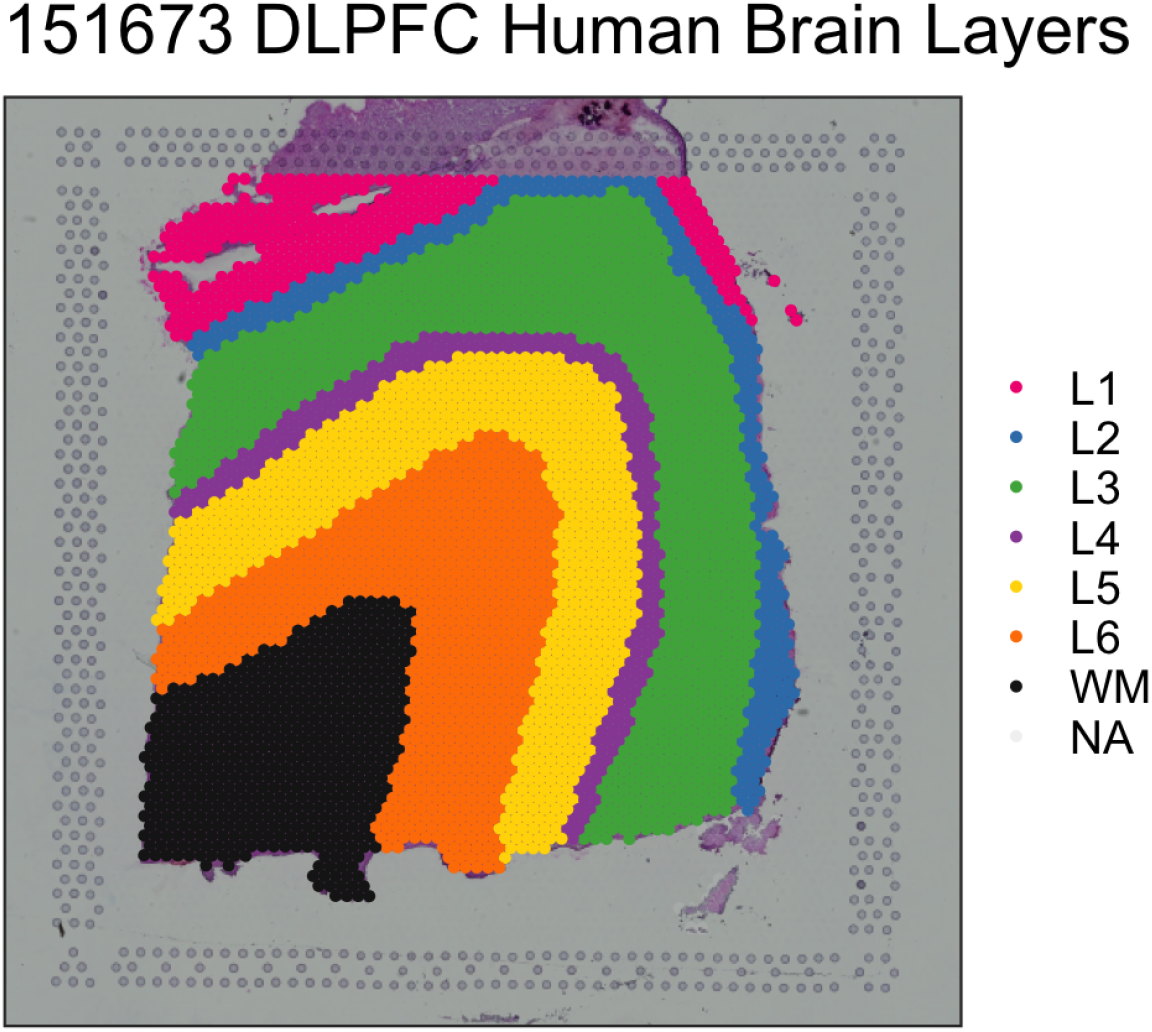
Simulation setting 1: The manual annotation layer of the DLPFC dataset. The spots span six cortical layers and white matter (WM), with the following distribution: L1 (273 spots), L2 (253), L3 (989), L4 (218), L5 (673), L6 (692), and WM (513).

**Fig. S3:**
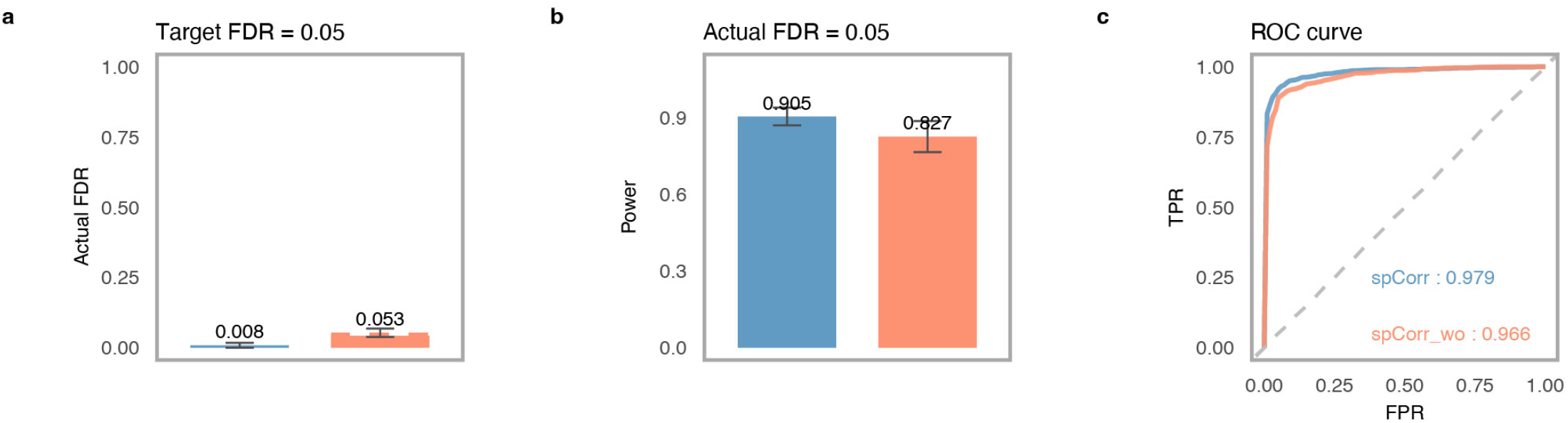
Simulation setting 1: Including covariates in marginal distribution modeling (step 1) improves FDR control, statistical power, and AUROC in spCorr. **a**. Actual FDRs achieved by spCorr with and without covariate adjustment at a target FDR of 0.05 (Benjamini–Hochberg adjusted *p* ≤ 0.05). Incorporating covariates yields better FDR control. **b**. Statistical power of spCorr with and without covariates, under the actual FDR = 0.05 cutoff. Modeling covariates improves detection power. **c**. ROC curves and corresponding AUROC values. The default spCorr (with covariates) achieves a higher AUROC than the spCorr (without covariates).

**Fig. S4:**
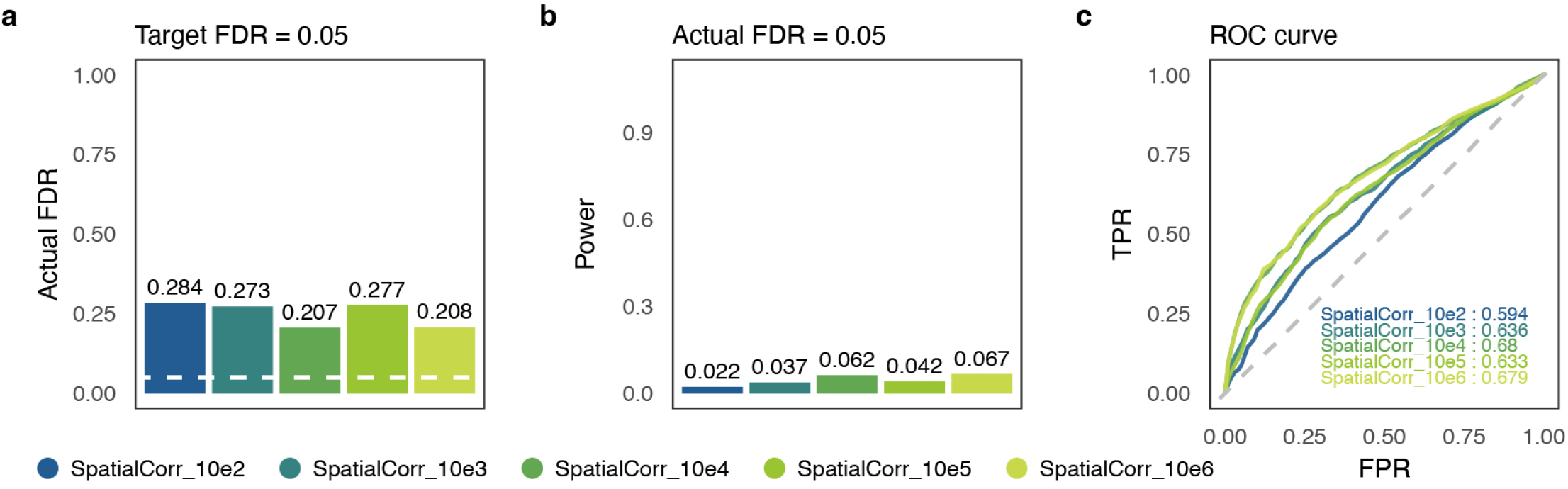
Simulation setting 1: Sensitivity analysis of the kernel bandwidth parameter *γ* in SpatialCorr. Performance was evaluated with *γ* ∈ {10^2^, 10^3^, 10^4^, 10^5^, 10^6^}. **a**. Actual FDR at a target FDR level of 0.05. **b**. Statistical power at the actual FDR level of 0.05. **c**. ROC curves with corresponding AUROC values. Results show that varying *γ* has minimal impact on FDR, power, or AUROC.

**Fig. S5:**
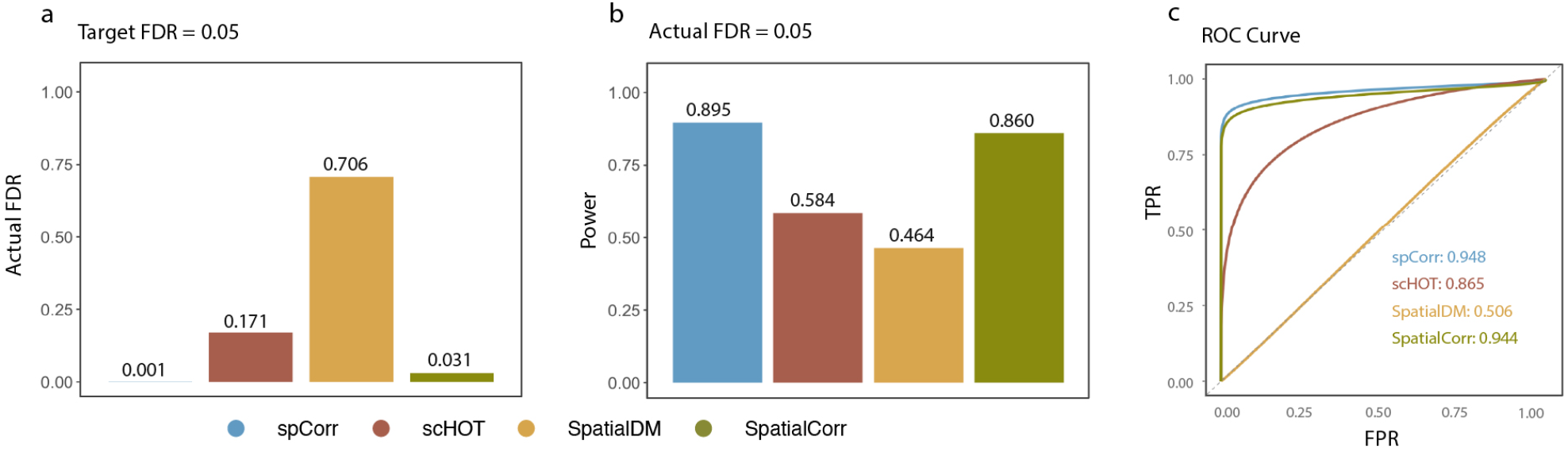
Simulation setting 2: Performance of spCorr, scHOT, SpatialDM, and SpatialCorr under a fully synthetic framework based on a bivariate negative binomial model with spatially varying copula structure. **a**. Actual FDR at a target FDR level of 0.05. **b**. Statistical power at the actual FDR level of 0.05. **c**. ROC curves with corresponding AUROC values. Results show that spCorr achieves the best FDR control, highest power, and largest AUROC.

**Fig. S6:**
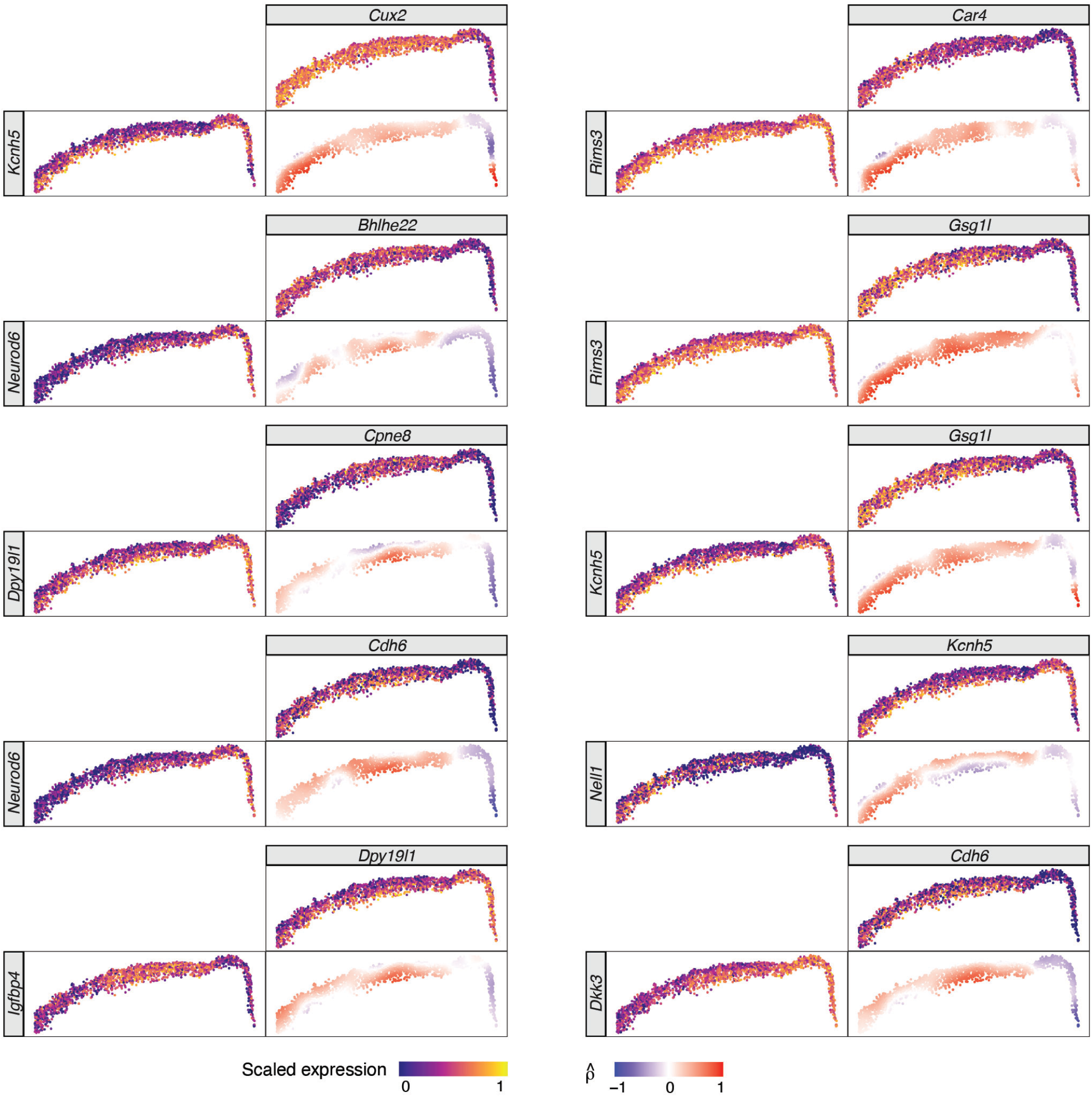
Additional visualization of spCorr-inferred local correlation estimates. Local correlation estimates for the top ten SVC gene pairs, together with their scaled gene expressions, are plotted in the original two-dimensional space.

**Fig. S7:**
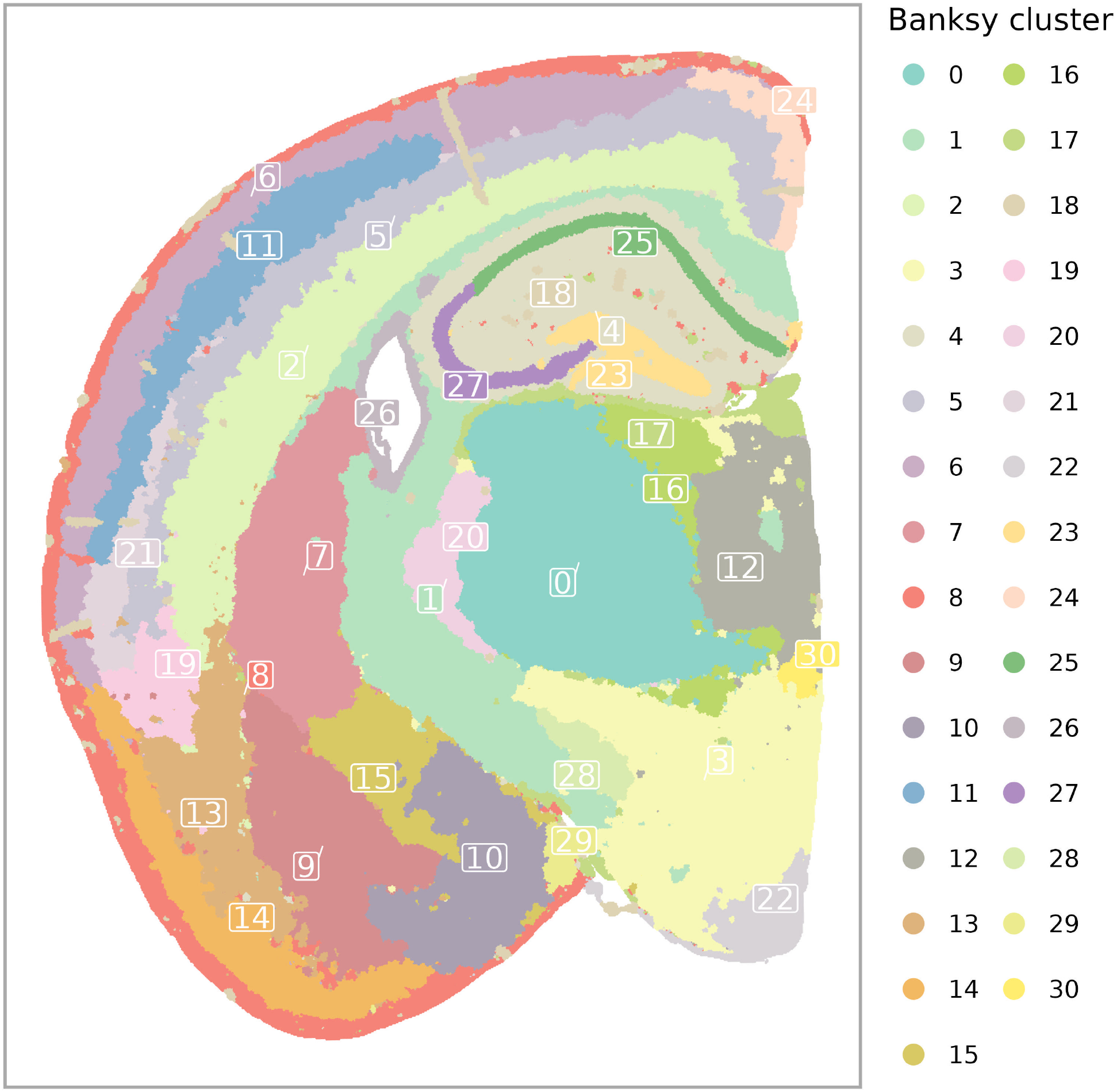
Spatial domains identified by BANKSY in mouse brain hippocampus Visium HD data. BANKSY segmentation results on the Visium HD dataset, showing 31 spatially-defined tissue domains derived using BANKSY with `lambda = 0.8`and `k_geom = 50`. Among them, domains 25 and 27 were selected as corresponding to the hippocampal pyramidal layer for analysis.

**Fig. S8:**
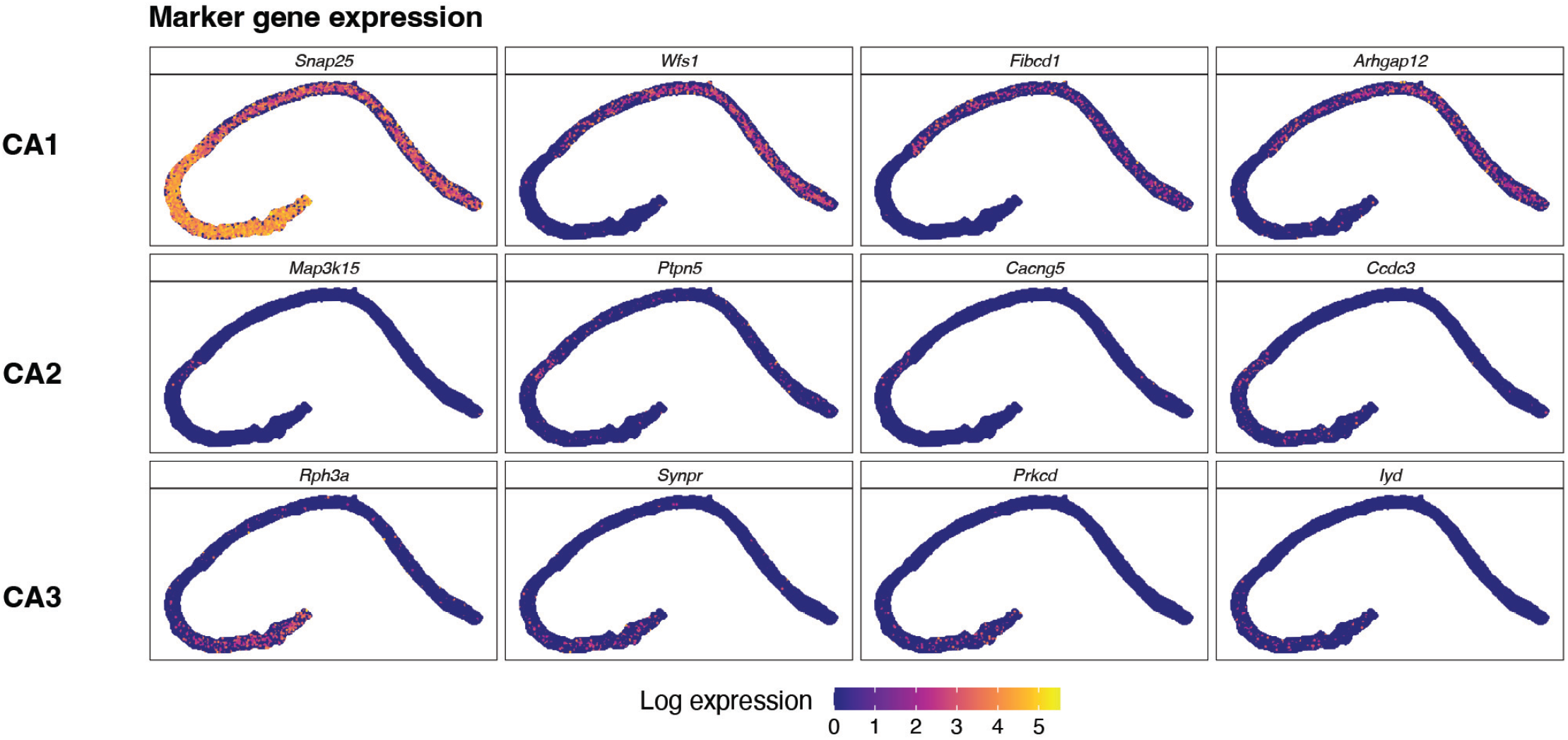
Marker gene expression for hippocampal domain annotation. Expressions of canonical marker genes are used to delineate the CA1, CA2, and CA3 domains within the hippocampal pyramidal layer.

**Fig. S9:**
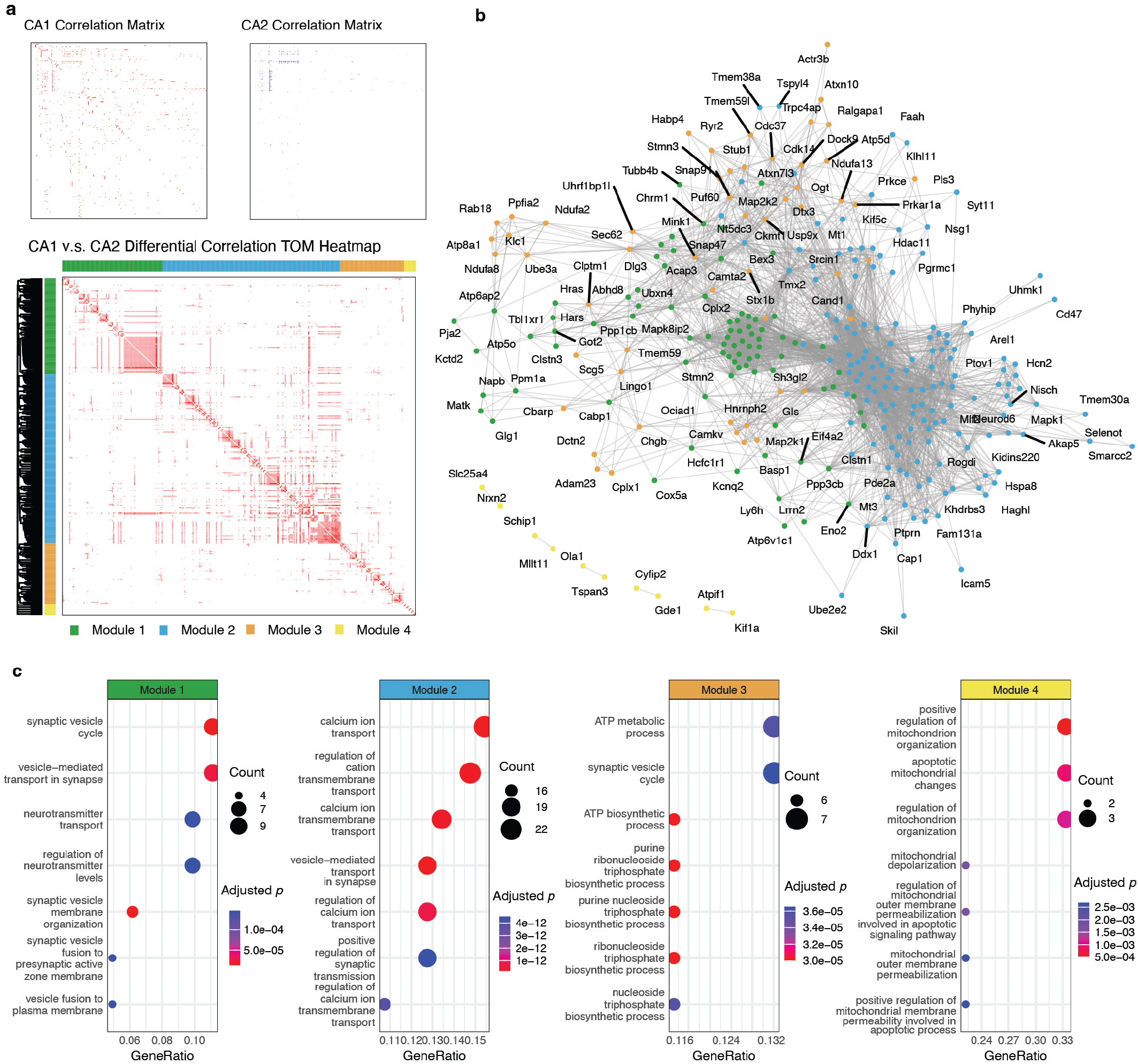
Differential correlation analysis without marginal distribution modeling leads to less functionally relevant gene modules. **a**. Correlation matrices for CA1 and CA2 domains, and the topological overlap matrix (TOM) heatmap of differential correlation between CA1 and CA2. Gene modules identified via hierarchical clustering are annotated by color. **b**. Gene correlation network constructed from significantly DC gene pairs, colored by module membership. **c**. GO enrichment analysis for each module. Although four modules are detected, the enriched biological processes are less clearly aligned with known functional distinctions between CA1 and CA2, illustrating the importance of adjusting for domain-specific expression differences during marginal distribution modeling (step 1 in spCorr).

**Fig. S10:**
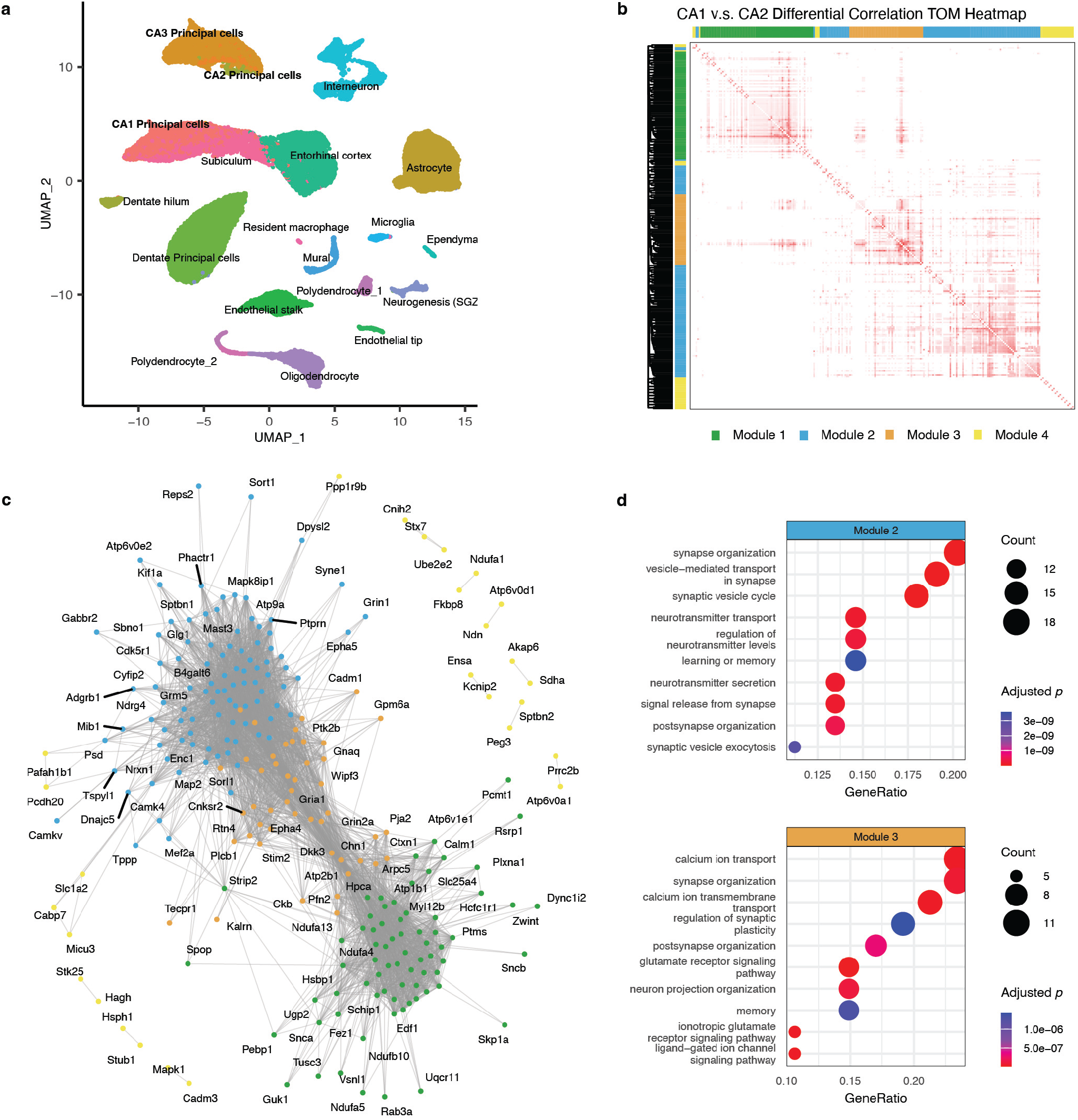
spCorr discovers biologically relevant correlation shift between cell types in scRNA-seq data. **a**. Cell type annotations for a mouse hippocampus scRNA-seq dataset, visualized via UMAP. Pyramidal neurons (CA1, CA2, and CA3 principal cells) were selected for downstream differential correlation analysis. **b**. TOM heatmap of the differential correlation matrix between CA1 and CA2 principal cells. WGCNA clustering identified four gene modules. **c**. DC network among significantly shifted gene pairs between CA1 and CA2. **d**. GO enrichment analysis for two representative modules from (c), highlighting biological functions related to synaptic plasticity and calcium ion transport.

